# Deep surveys of transcriptional modules with Massive Associative K-biclustering (MAK)

**DOI:** 10.1101/2022.08.26.505372

**Authors:** Marcin P. Joachimiak, Cathy Tuglus, Rauf Salamzade, Mark van der Laan, Adam P. Arkin

## Abstract

Biclustering can reveal functional patterns in common biological data such as gene expression. Biclusters are ordered submatrices of a larger matrix that represent coherent data patterns. A critical requirement for biclusters is high coherence across a subset of columns, where coherence is defined as a fit to a mathematical model of similarity or correlation. Biclustering, though powerful, is NP-hard, and existing biclustering methods implement a wide variety of approximations to achieve tractable solutions for real world datasets. High bicluster coherence becomes more computationally expensive to achieve with high dimensional data, due to the search space size and because the number, size, and overlap of biclusters tends to increase. This complicates an already difficult problem and leads existing methods to find smaller, less coherent biclusters.

Our unsupervised Massive Associative K-biclustering (MAK) approach corrects this size bias while preserving high bicluster coherence both on simulated datasets with known ground truth and on real world data without, where we apply a new measure to evaluate biclustering. Moreover, MAK jointly maximizes bicluster coherence with biological enrichment and finds the most enriched biological functions. Another long-standing problem with these methods is the overwhelming data signal related to ribosomal functions and protein production, which can drown out signals for less common but therefore more interesting functions. MAK reports the second-most enriched non-protein production functions, with higher bicluster coherence and arrayed across a large number of biclusters, demonstrating its ability to alleviate this biological bias and thus reflect the mediation of multiple biological processes rather than recruitment of processes to a small number of major cell activities. Finally, compared to the union of results from 11 top biclustering methods, MAK finds 21 novel *S. cerevisiae* biclusters. MAK can generate high quality biclusters in large biological datasets, including simultaneous integration of up to four distinct biological data types.

**Author summary:** Biclustering can reveal functional patterns in common biological data such as gene expression. A critical requirement for biclusters is high coherence across a subset of columns, where coherence is defined as a fit to a mathematical model of similarity or correlation. Biclustering, though powerful, is NP-hard, and existing biclustering methods implement a wide variety of approximations to achieve tractable solutions for real world datasets. This complicates an already difficult problem and leads existing biclustering methods to find smaller and less coherent biclusters. Using the MAK methodology we can correct the bicluster size bias while preserving high bicluster coherence on simulated datasets with known ground truth as well as real world datasets, where we apply a new data driven bicluster set score. MAK jointly maximizes bicluster coherence with biological enrichment and finds more enriched biological functions, including other than protein production. These functions are arrayed across a large number of MAK biclusters, demonstrating ability to alleviate this biological bias and reflect the mediation of multiple biological processes rather than recruitment of processes to a small number of major cell activities. MAK can generate high quality biclusters in large biological datasets, including simultaneous integration of up to four distinct biological data types.

## Introduction

Innovations in scaling measurement technology based on sequencing, mass-spectrometry, and cytometry are driving the development of methods that can efficiently extract statistically supported predictions of gene function, interaction, and contribution to phenotype. One of the first critical steps of such analyses is the detection of patterns in the data which group together features across measured molecules or bioentities that are similar over a subset of conditions. This reduces the dimensionality of the data and provides first hypotheses about associations among these features.

### State of the art in biclustering

A number of different types of algorithms have emerged to perform this type of pattern detection and data reduction on two-dimensional data including hierarchical (1), K-means (2), clustering methods (3, 4); random forests (5), Bayesian networks (6), support vector machines (7), singular value decomposition (8) and principal component analysis (9); and biclustering (10). Biclustering methods have emerged as one of the most popular methodologies (10–14), because they are able to find local, context-specific patterns, are better at identifying signals in large, noisy datasets with overlapping patterns, and can include information from both data axes as well as additional data. In biology, most biclustering methods have been applied to microarray data (10-12, 14-19) but also to other data types such as metabolite levels (20), drug interactions (21–23), RNA multiple sequence alignment (24), phenotype data (25), protein-protein interaction mass spectrometry data (26–28), and as part of a machine learning pipeline to identify literature associations (29). Examples of impactful discoveries from these algorithms include functional assignments for uncharacterized genes (15), identification of transcriptional modules (30), identification of transcriptional modules with putative transcription factor (TF) binding sites and support in other data types such as protein interactions, pathway membership, phylogenetic profiles, operon associations, and sequence motifs (13, 31), breast cancer classification and prognosis (32), identification of associations between transcriptional modules and environments suitable for predicting microbial response to environmental change (33), and large scale biomedical relationships derived from literature (29).

While biclustering can be performed on many different types of data, here we focus on differential gene expression as an example of large-scale experimental measurements. This example is analogous to other common biological data types such as taxonomic and functional abundance profiles in metagenome data, cell counts and quantitative cell phenotypes, cell line responses to chemical treatments, and other data generated with high throughput technologies. These experimental data sets are usually composed of a collection of experiments each containing a set of conditions such as a time series or field sampling, where each condition may be the summary of a set of replicate samples.

Traditional clustering techniques, such as hierarchical clustering, measure similarity globally across all data points, such as individual gene expression conditions, in contrast to a local comparison, which is able to only consider a subset of the data (e.g. only conditions exhibiting a certain data pattern). Global comparisons may be unable to identify local patterns such as from condition-specific transcriptional regulation, e.g. in response to a specific environmental cue like heat shock, due to the lack of signal in the remainder of the conditions not involved in that biological response or overwhelming of the local signal with data from the other conditions (34, 35). Global comparison approaches, like hierarchical clustering or K-means, have additional limitations, such as sensitivity to noise and inability to identify overlapping clusters (36). Biclustering methods seek to overcome these limitations by identifying subsets of columns (e.g. conditions in gene expression data) over which rows (e.g. genes) show similar behaviors. In the following sections we also compare two-dimensional hierarchical clustering (2D-HCL) approaches against biclustering methods and find that they perform worse than other methods. For a given gene expression dataset, an ideal biclustering algorithm should identify all the condition-specific gene co-expression biclusters exhibiting coherent patterns across a subset of conditions. Coherent patterns are identified when trends in the data can be closely fit to a mathematical model (14), such as constant expression or constant column expression.

For discovery in gene expression data we consider a subset of possible bicluster pattern types, namely constant, constant row, constant column, and checker or composite row and column patterns. Constant biclusters represent constant expression across a set of genes and conditions, while constant row and constant column biclusters represent constant expression of each gene in a set across a series of conditions, with different values for each gene, or constant expression of a set of genes in a series of conditions, with different values for each condition. The identified biclusters should i) only contain genes and conditions conforming to the overall bicluster pattern thus having a low false positive rate and high precision, ii) contain all relevant genes and conditions conforming to the pattern, i.e., high true positive rate and high recall, iii) exhibit high coverage of significant patterns in the data set, and iv) capture manifestations of biological mechanisms in the data that exhibit coherent data patterns, for example biclusters in gene expression data should be statistically enriched for functional categories and transcriptional regulation corresponding to coordinated control of gene expression.

### Challenges and limitations of extant biclustering algorithms

However, biclustering can scale poorly and in fact, it has been shown that even simple variants of a search for all biclusters in a dataset is an NP-hard problem (10), placing a strong limitation on the size of the data sets for which exact solutions are available. In practice however, exact solutions for popular clustering methods can also be NP-hard (e.g. K-means (37)) or scale poorly (e.g. at least O(N^2^) for hierarchical clustering (38)). Thus in general, tractable methods need to strike a balance between computational complexity, accuracy, and coverage of the significant patterns in the data. Such a balance is achieved by implementing approximations that are reflected in algorithmic designs and parameters such as the number of desired biclusters, their degree of overlap, or their size. However, these quantities can be difficult to assess in biological data without ground truth, rendering parameter tuning and exploration a challenging problem in of itself and contributing to the lack of reliable unsupervised methods. With the rise of accessible high-performance computing (HPC), there is an opportunity to improve both performance and scalability, while decreasing the need for supervision.

Available biclustering methods have proven useful for functional discovery and regulatory network inference but they are also limited by a number of key elements that have prevented their widespread adoption. These issues span operational aspects such as requiring many parameters and finding an optimal parameter set, as well as poor support of biclustering results in terms of post-analysis and visualizations, partly due to the complexity of the data and biclustering results. The existing issues also involve aspects of the biclusters themselves that may hinder uncovering biological relationships such as preprocessing the input data to reduce dataset size (e.g., removing rows or columns with low variance) or complexity (e.g., converting to integer or binary values), introducing bicluster sizes biases, limits or biases in exploration of the bicluster search space such as truncation of searches, as well as restrictions on the standard bicluster pattern type models. Formal evaluations of biclustering results are critical for revealing these issues and comparing and contrasting method performance.

Previous biclustering method evaluations (36, 39, 40) have shown that biclustering methods often yield biclusters that are too small, i.e. genes or conditions with a highly similar pattern are erroneously omitted (false negatives). One example of bias towards smaller biclusters is the popular mean squared error (MSE) (41) measure of gene expression bicluster coherence. The MSE is an important measure since it makes no assumptions about the data, can be applied to different data patterns, and has a biological interpretation, namely on average, how similarly a set of genes responds to a set of experimental conditions. However, the MSE can be trivially minimized by shrinking the size of a bicluster (42), inevitably resulting in false negatives and potentially patterns too small to validate with biological term enrichment analysis. Furthermore, biclusters can be too large or too small, too few or too many, with genes which are not significantly differentially expressed, or offering low coverage of the interesting values in the data (e.g. significant differential gene expression). In addition, the nuances of input data sets can produce a number of special cases including missing values and global versus local data patterns, which have the potential to impact the results (e.g. data patterns may not be recognized due to a lack of imputation for missing data or due to data pre-processing such as removing features below minimum expression thresholds (13) or avoiding sets of genes or conditions with smaller average changes in expression (30)). Finally, most biclustering algorithms cannot operate simultaneously on multimodal data with different data types and cannot therefore evaluate and exploit the complementarity therein. The need to interrogate multiple layers of data types has driven the development of new methods [2,17-20], though these still suffer from other limitations mentioned above and are even more difficult to evaluate with simulated and real-world data.

### Novelty of the Massive Associative K-biclustering (MAK) approach

To address a number of the existing biclustering challenges, we developed a biclustering algorithm called Massive Associative K-biclustering (MAK). MAK is an associative method because it associates groups of objects by identifying shared data patterns within individual or multimodal data of different types. The K in K-biclustering represents the number of data types being simultaneously biclustered. At the core of the MAK algorithm are a series of innovations aimed at improving discovery: 1) adjusting for bicluster score size biases, 2) performing step-wise optimal bicluster search moves, 3) performing many parallel searches in an HPC environment, and 4) providing a final nonredundant set of biclusters across different pattern types, while preserving a desired degree of overlap. We implemented statistical bicluster coherence criteria to improve measures of bicluster coherence for a variety of bicluster pattern types and canonical data types. We adjust these criteria via a statistical biclustering innovation by using dataset-specific null models. The null models allow scoring of biclusters in a data adaptive manner by adjusting for statistical criteria biases with respect to bicluster size and allowing rigorous combinations of different criteria within and across bicluster pattern types, as well as input data types.

In this study we use gene expression data as an example of continuous numerical data; however, we also introduce scoring criteria spanning three other common biological data archetypes: pairwise interactions, ranked features, and binary features. We demonstrate MAK biclustering with different data type combinations and evaluate these results using other biclustering methods. MAK is a greedy algorithm but performs stepwise optimal bicluster searches by testing all possible moves for the move type chosen at each step. Overall, MAK uses three types of approximations: 1) the null distributions as smoothed representations of bicluster coherence values computed for randomly sampled biclusters of specific sizes, 2) bicluster search space bounding by the limits of the computed null distributions for bicluster size ranges, and 3) MAK pseudo-random walks using a fixed sequence of move attempts, which is repeated until stopping conditions are met. We have tested the MAK HPC implementation on datasets of up to 100,000 rows and 10,000 columns, using default parameters including stepwise optimality. Thus, MAK can provide an important benchmark for comparing other biclustering methods. To this end we perform a multi-faceted biclustering comparison on a real world dataset, in addition to simulated data comparisons, across a set of methods that have been designed for real valued matrix input data and demonstrated to have top performance on gene expression data (17, 18, 31, 43–49).

### Multi-faceted biclustering method evaluation

Another challenge is the systematic evaluation of biclustering methods. There are two distinct approaches to their evaluation: one is designing simulated datasets with known implanted biclusters, and the second is to assess performance on real biological data where the true answers are effectively unknown. The former aims to statistically assess the performance of a method, allowing measurement of standard metrics like true and false positive rates, while the latter is instrumental for biological validation. Currently, simulated data testing cannot replace biological validation. However, the size and complexity of real-world datasets impose limitations on testing procedures and cannot provide statistical details on performance due to a lack of gold standards. A number of systematic biclustering evaluations have considered summaries of statistical enrichments of function terms (36, 39, 50) and in rarer cases also enrichments for known regulation (31). However, attempts to systematically validate the scoring of biclusters and to relate them to biological mechanisms are lacking. The one exception has been the cMonkey method (13), where the authors identified gene expression biclusters composed of genes sharing putative regulatory binding sites and subsequently applied this to infer and predict transcriptional responses (33).

To understand biclustering method performance it is necessary to consider both experimental and realistically simulated data, as ground truth for biological data is in general not available. We performed two different synthetic data evaluations to demonstrate the efficacy of MAK. First, we evaluated 94 different MAK bicluster coherence criteria using a high throughput bicluster coherence criteria test-bed targeting the basic bicluster pattern types implemented in MAK. This simulated data was designed to cover standard biclustering test scenarios involving variations in the size, overlap, orientation, and coherence of biclusters, but also to be easier to interpret, more compact, and more realistic with respect to a gene expression compendium. In the high throughput criteria evaluation, we use the F1 measure (51) to assess biclustering methods, which represents the analogy to binary classification where bicluster genes and conditions belong either inside or outside the implanted pattern. The F1 measure is a single statistic that represents the harmonic mean of precision and recall, with high values corresponding to low false positive and high true positive rates. Evaluation on the simulated data enabled the selection of optimal MAK bicluster criteria for the algorithm implementation, which we subsequently evaluated against other methods using a larger and broader biclustering evaluation framework (52), which has been reused and exhibited consistency in results from different groups (44). We demonstrate that on this simulated data MAK is only outperformed on certain pattern types by two recent methods (EBIC [41], RecBic [42]) and exhibits state of the art performance while having the lowest variance for each bicluster pattern type.

On real-world data the resultant biclusters are open to biological interpretation. For example, constant and column gene expression biclusters indicate genes with coordinated regulation across a set of conditions. In these cases for the gene set associated with the bicluster one might expect a consonant enrichment in TF binding sites and common functional themes. Row biclusters have a similar interpretation but whereas non-constant column biclusters might show how expression of a set of genes changes together in multiple conditions, row biclusters show which genes maintain relatively constant expression across a set of conditions. Finally, the composite checker pattern biclusters contain both a pattern and an anti-pattern, indicating two gene sets, the first being genes with opposite expression in different conditions and a second gene set with an opposite pattern.

To evaluate MAK biclustering against other methods on real-world data we use a well-studied yeast gene expression dataset (13, 53). Most biclustering evaluations on real biological data have relied exclusively on Gene Ontology (GO) (54) term enrichment. We attempt to circumvent limitations stemming from biases and incompleteness of individual biological term enrichments, by allowing for multiple types of statistical enrichments per bicluster as combinations of GO terms, KEGG pathways (55), TIGRFAM roles (56), and curated regulatory associations (57). Unfortunately functional enrichment is limited by our current understanding and representation of biological functions and our ability to accurately and completely annotate biological data. For example, unrecognized facets of traditionally static cellular processes such as specialized ribosomes and protein expression are being reported (58, 59). Beyond functional annotations, experimentally determined regulatory associations are rare and only available for model organisms (60–62) and even these are expected to be incomplete, rendering thorough biological evaluation of biclustering results difficult. This is even more challenging for condition-dependent information since it is not available for most experimental data, such as TF binding or protein interactions. Thus, to provide additional context beyond standard biological term enrichment, we also introduce a data-driven bicluster set quality validation approach based on easily computable bicluster utility criteria applicable to real world datasets without a gold standard reference. The method comparisons also involve bicluster size distributions, the relationship between enrichments and bicluster coherence, enriched GO term combinations, and overlap between bicluster result sets.

We demonstrate the biological efficacy of MAK and its data type integration capabilities through biclustering yeast gene expression data, as well as combinations of expression data with phylogenetic profiles derived from related genome ortholog conservation patterns, and experimentally determined TF binding associations and protein-protein interactions. MAK maximizes the joint distribution of bicluster coherence and biological enrichments and showed maximal performance when multiple data types were included. On gene expression data alone, MAK found 21 novel biclusters with enriched biological features, which were not found amongst the 1655 biclusters from 11 other top biclustering methods. MAK results, when including experimental TF-gene associations, exhibited the second most unique TF enrichments, second only to cMonkey, which relied on sequence motif detection and multiple data types and had low bicluster coherence and contrast. Finally, MAK’s results when applied to gene expression and three additional data types showed the greatest joint performance for bicluster coherence and total biological enrichments, indicating there is additional information beyond gene expression and that MAK can utilize this information to find the most coherent patterns in data.

## Results

### MAK method overview

The MAK framework consists of modular components. These include the core MAK algorithm as well as a suite of bicluster scoring criteria for different data types along with functionality to generate the null models. The framework also consists of a simulated data test-bed to evaluate biclustering algorithms and scoring criteria performance. Included is a collection of methods to execute bicluster discovery strategies in HPC environments. Finally, a collection of tools targets post-analysis of bicluster sets including assessment of features of biclusters from arbitrary datasets.

### MAK bicluster scoring criteria

The overall MAK bicluster coherence scoring criterion is a summation of data type- and pattern-specific criteria. Each criterion is based on statistical functions along with data-specific null models. The null models convert bicluster criteria into scores based on how extreme the value is compared to a distribution of bicluster criteria values recorded from a population of randomly sampled data submatrices of specific sizes (see Methods). The null distributions provide dataset-specific context for bicluster criteria and need to be computed for each combination of input dataset and scoring criterion targeting a specific bicluster pattern type but represent only a few percent of the total MAK runtime on yeast gene expression data. The null models allow adjustment for bicluster size biases, such as the bias of the MSE for smaller biclusters, and to accommodate variations among the distributions of the different criteria. We use the latter property to rigorously combine multiple bicluster criteria for the same data type as well as different criteria for different data types.

Biclusters can have three types of basic patterns: constant, column, and row (14), and we also consider a checker pattern composed of these three (Figure 1). Row and column patterns, where rows are genes and columns are conditions in gene expression data, can be further classified by the mathematical model describing their relationships, such as constant, additive, or multiplicative (14). In additive and multiplicative row or column biclusters, the row- or column-wise values differ from each other by an additive or multiplicative constant. Additive relationships combined with multiplicative is another possibility, referred to as shift-scale (14). Checker biclusters, in data with signed values, correspond to two column or row patterns stacked in opposite, and their coherence can be measured with row and column criteria after taking the absolute value of the input data.

**Figure 1.**
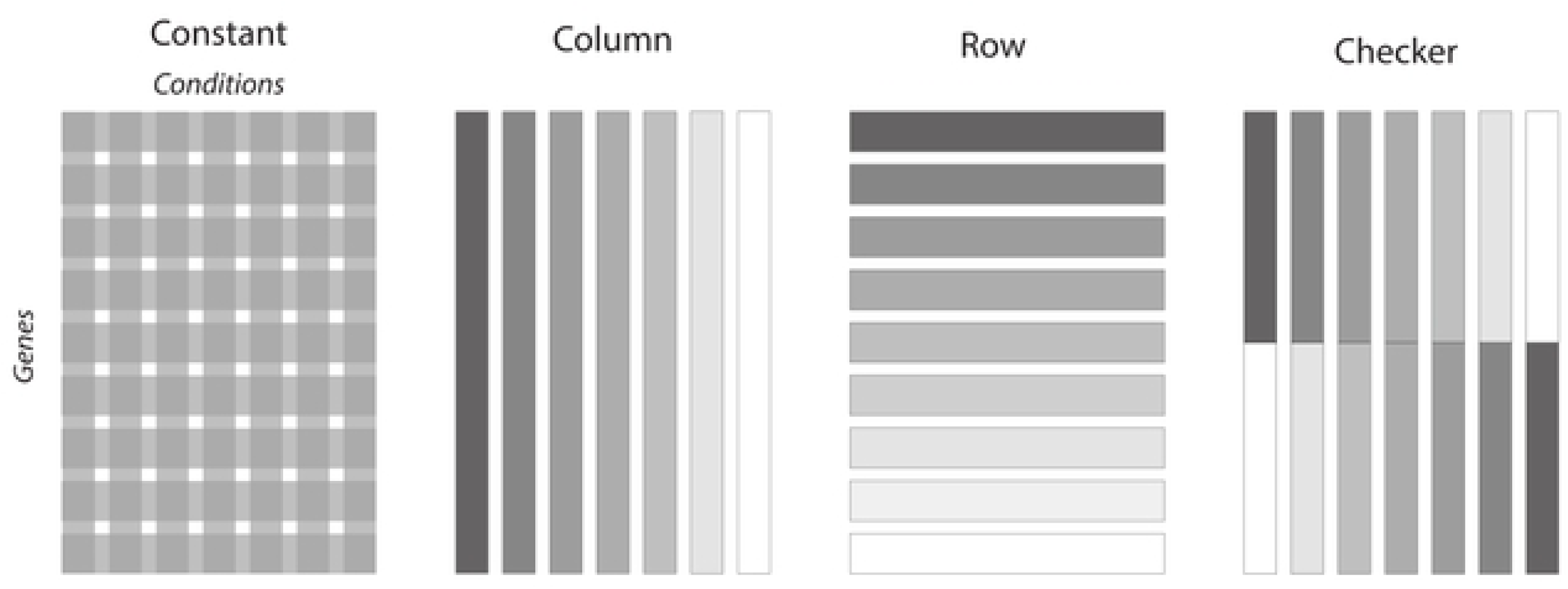
Basic coherent bicluster data patterns. Cartoons of idealized bicluster patterns. A constant pattern is equivalent with respect to coherence measured by rows or columns. On the example of gene expression data, constant row coherence corresponds to a set of genes with constant expression across a series of conditions, with the constant value differing between genes. Conversely, constant column coherence corresponds to a set of genes with constant expression in each condition, but with the constant value differing between conditions. A checker pattern corresponds to two row or column patterns arranged together vertically or horizontally but in opposite orientations.

We develop scoring criteria that evaluate a size-corrected coherence of bicluster rows, columns, or both. The scoring criteria favor biclusters with elements (rows or columns), which are positively and, for signed value data, optionally also negatively associated. The null models ensure that the criteria are evaluated fairly against the data and facilitate combinations of multiple criteria. Specifically, we compute a probability that a randomly sampled bicluster of the same size would have a criterion value greater than the bicluster resulting from the current move, parameterized with the Cauchy distribution to better account for extreme values. Thus when evaluating moves in the MAK algorithm, we can score the objective by how much better the current bicluster is compared to the expectation from random chance.

The MAK bicluster criteria suite (Sup. Info.) includes modifications of well-known criteria like MSE, in the form of column (MSEC) or row-based (MSER) mean squared error. These variants of the MSE represent the mean deviation from the row means or columns means (represented as a relative ratio to the total MSE, see Methods), and correspond to mean row or column variability, respectively. Additional top ranking bicluster criteria were Kendall’s W (63) non-parametric criterion which measures similarities in samples across multiple test attempts, as well as a regression criterion based on a fixed effects model (FEM) (64), which measures the row (FEMR) or column (FEMC) variability within the block versus in the remaining data. In all, we evaluated 94 different bicluster scoring criteria and their combinations, including criteria that have been traditionally used in gene expression clustering analysis (i.e., Pearson correlation, Euclidean distance).

In addition to the bicluster criteria measuring gene expression patterns, we developed criteria that score agreement of ranked TF binding preferences for a set of genes (based on protein binding microarray data (65)), binary pairwise interactions (i.e., protein-protein interactions (PPI) (66)), and binary features (i.e., gene phylogenetic profiles across yeast genomes (67)) (see Methods). These criteria serve as demonstrations of integrating rank, interaction, and feature data archetypes, and therefore are more generally applicable. For example, the MAK gene expression criteria can be directly applied to quantitative phenotypes, the TF criterion can be applied to gene rank data, the interaction criterion to genetic interactions, and the gene feature criterion to gene functional annotations, and similarly for data where the primary entity is other than genes, such as taxa or chemicals.

The sole requirement is that the y-axis of the primary data matrix, here gene expression data, matches the y-axis in the other data types (and also the x-axis in the case of interactions). Modifications of these four canonical data type criteria as well as introduction of novel criteria are supported within the MAK framework.

### MAK algorithm

MAK includes a customizable algorithmic framework allowing addition and removal of data types and bicluster criteria, specification of how criteria and data types are combined, as well as modification of individual bicluster searches including the overall discovery strategy. The MAK algorithm can be embedded at different stages in a bicluster discovery strategy, supported by the MAK framework and tools. Figure 2 outlines the details of the MAK algorithm (Figure 2.A) and the MAK bicluster discovery strategy used for results in this study (Figure 2.B). MAK begins by identifying a set of starting clusters or biclusters for the input data. The starting points can be chosen, for example, as we do here using 2D-HCL and a bicluster size range restriction (see Methods) (Figure 2.B), which represents a complete tiling of the data. MAK can use any gene or condition clusters or sets, as well as random or candidate biclusters as starting points, as well as combinations of thereof. For each of these starting points MAK chooses to add one or more rows (e.g. genes) or columns (e.g. conditions) if they improve the score (Figure 2.A). The full series of moves for a single starting point corresponds to one MAK bicluster search trajectory, and the endpoints of these trajectories are considered candidate biclusters subject to post-processing operations such as pooling, thresholding, and non-redundancy selection or merging.

**Figure 2.**
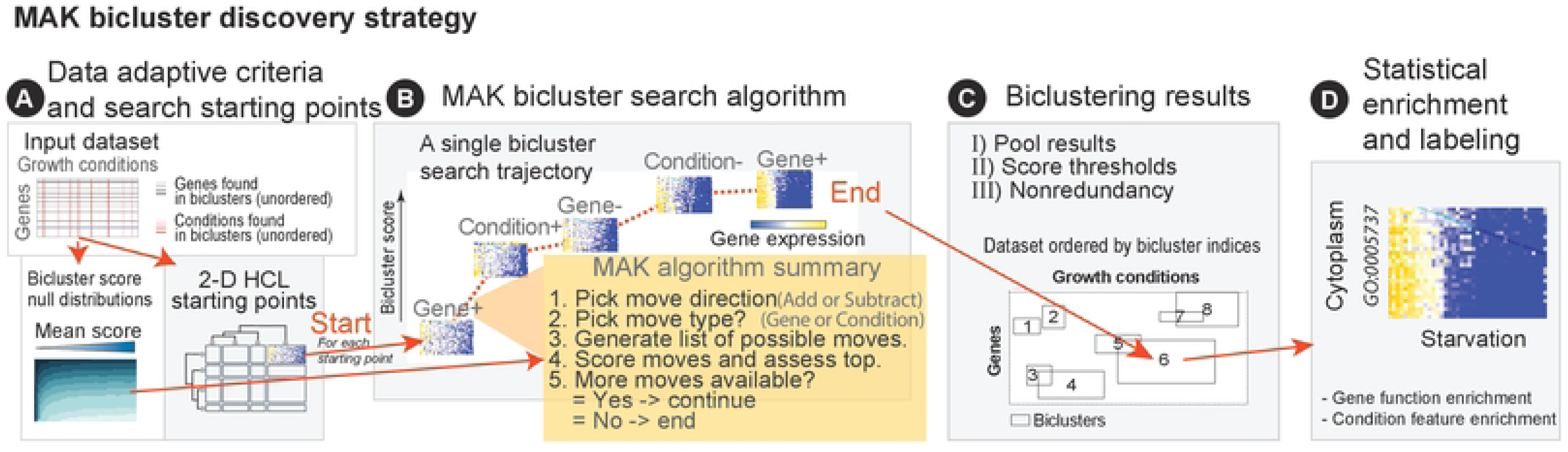
MAK algorithm and bicluster discovery strategy. A schematic of the MAK algorithm (**A**) and MAK bicluster discovery strategy (**B**). The MAK algorithm is based on a search of bicluster space and the starting points in the biclustering discovery strategy are generated by 2D-HCL. The search consists of choosing a move direction (addition or subtraction), a move axis (e.g. gene or conditions for gene expression data), and a move type (single or batch). The underlined algorithmic features are available optimizations that allow reduction of the computational cost. The bicluster discovery strategy detailed in 1.B outlines the main stages starting from an input dataset, generating null distributions and starting points, running multiple rounds of MAK biclustering, and arriving at a set of the final biclusters after filtering by score, nonredundancy, and pooling results from different bicluster patterns.

The MAK bicluster scoring criteria can be combined to create a single objective function for each move. To do this, the user first selects a subset of bicluster coherence criteria to create the objective function, where the criteria may be for the same or different data types. Here we report results for the combination of column MSE, Kendall’s W criteria, and FEM, which were selected based on a formal evaluation on simulated data (Sup. Info.). The data-dependent null model is applied to each raw criterion, and the resulting values are summed for a single data layer or weighted and summed across data layers into a single score.

MAK can explore all bicluster search space moves, as determined by the move sequence, and therefore can select the move that best improves the bicluster score at each step, hence achieving stepwise optimality. This stepwise process is repeated until there are no more moves available to further improve the current bicluster score. Thus, MAK is a greedy, stepwise optimal algorithm with heuristics to guide the random walk and limit its length. The algorithm relies on a directed pseudo-random walk, based on a repeating move sequence starting with randomly picking gene addition or gene deletion. For background, the MAK algorithm has the concept of move modes and move types. MAK utilizes a move mode sequence, such as batch moves followed by singles moves, and for each move mode executes a repeated move type sequence of gene or experiment addition or deletion (Sup. Info.). Each gene move is followed by the corresponding condition addition or deletion move, and vice versa following a condition move. Thus, the directed random walk consists of repeated sets of moves corresponding to two allowed move sequences: (g- -> e- -> g+ -> e+) and (g+ -> e+ -> g- -> e-). This heuristic strives to maintain a balance of moves and to allow the bicluster pattern to stabilize across both axes of the data (e.g. testing other moves after a gene addition before another gene addition). If in the last move mode (e.g. single moves in a (batch, single) MAK mode run) all four possible move types have been tried and the last move type does not lead to a higher score, then the algorithm stops and a final block, a candidate bicluster, is reported. Additionally, moves may not be possible due to size range limits on the expected block size or due to null distribution bounds, and these cases are treated as attempted moves and contribute to triggering stopping criteria.

Multiple rounds of MAK bicluster discovery can involve large numbers of searches and computation; hence optimizing starting point size, null distribution bicluster size ranges, the move sequence, and stopping heuristics are desirable. For this work we ran MAK trajectories for all starting points as a complete reference, however trivial optimizations are available to significantly reduce computation in this step (Sup. Info.).

In the MAK bicluster discovery strategy, multiple rounds of bicluster searches are used to discover overlapping biclusters or ones with comparable coherence. To combine information from many MAK bicluster searches we perform a nonredundant selection of overlapping biclusters by choosing representative biclusters with the highest score (see Methods). This selection can be performed across results from different bicluster pattern types as we show here for gene expression data; the selection can also be applied to results from different input data combinations. A cosine similarity overlap threshold heuristic, set at 0.25 or a quarter of the bicluster areas assuming equal sizes, attempts to remove redundancy while allowing for biological effects such as combinatorial regulation and preserving the structure of the biclusters. The overlap results may be organism and dataset-specific and the distribution of discovered bicluster overlaps can be used as guidance to tune thresholds as necessary. The overlap threshold can also be determined using heuristics, for example based on the lowest scores of biclusters with biological enrichments or by minimizing the number of biclusters, while maximizing their gene expression, and maintaining a reasonable bicluster size range.

MAK divides the bicluster discovery problem into many parallel computations, each computing an independent bicluster search trajectory. The default MAK configuration uses 2D-HCL starting points and performs batch moves followed by single moves. This MAK configuration scales as *O*(*N*^2^ + *M*^2^), where N is the number of genes and M the number of conditions. Without 2D-HCL or batch moves the complexity of the MAK algorithm is *O*(*N* + *M*), although in practice both choices are likely to improve convergence of bicluster search trajectories, hence reducing run time. For the yeast gene expression dataset with 6160 genes and 667 conditions, the average MAK trajectory from the four bicluster pattern types corresponding to the exemplar results completed in two hours and ranged from an average of 50 min. (column pattern) to almost 5 hours (row pattern). Note that the total CPU time for the MAK bicluster discovery pipeline is the wall time from starting the pipeline to obtaining the final output and includes both single CPU and CPU cluster wall time (S1 Table). For further details of the MAK framework see Sup. Info.

### High throughput simulated data MAK bicluster criteria evaluation and UniBic simulated data evaluation with other biclustering methods

We generated smaller, realistic simulated data to perform a high throughput evaluation of potential MAK bicluster scoring criteria across constant, column, row, and overlapping as well as non-overlapping bicluster pattern types (see Sup. Info.). Using the top identified MAK criteria, we compared MAK performance with other biclustering methods using the UniBic simulated dataset (52) (Figure 3). This evaluation has been reused by others, notably the EBIC biclustering algorithm that outperformed UniBic (44). We matched criteria for MAK pattern types to UniBic simulated bicluster pattern types (Figure 3C, Sup. Info.) and set parameters identical to those used for the high throughput MAK bicluster scoring criteria evaluation as well as the real-world yeast gene expression data biclustering, with starting point sizes and null bounds scaled to the input dataset size (Sup. Info.). The UniBic evaluation reports the relevance and recovery only for genes in each bicluster. Relevance measures the maximum overlap of biclusters with the ground truth and recovery measures the maximum overlap for each ground truth bicluster. Firstly, EBIC shows nearly perfect mean performance on this dataset although with considerable variance for patterns IV and VI (see (44)). We note that a more recent method, RecBic, used an analogous but larger simulated dataset and exhibited similar performance to EBIC (45). Across the six types of bicluster data patterns in the UniBic evaluation, MAK had the second best mean performance overall, after EBIC. MAK outperformed UniBic on 4 out of 6 pattern types (II, IV, V, VI), and tied for types I and III. In addition, the MAK results had among the lowest variance across all pattern types, indicating consistent performance. Pattern types II and III in the UniBic data represent column and row biclusters respectively, however these are devoid of noise and hence differ from the corresponding patterns in the MAK high throughput simulated, which modeled noise based on a real-world dataset. Thus for the type III row pattern a MAK row-wise Pearson correlation criterion outperformed the default MAK row criterion Kendall_FEM, which itself outperformed all but three methods (Sup. Info.). We also assessed MAK on the overlapping constant bicluster dataset from UniBic and found that across the four cases MAK was very close to the top method for the average of relevance and recovery across all cases (UniBic 0.775 vs. MAK 0.772, EBIC showed similar relevance but higher recovery (see (44)) (S2 Figure, Sup. Info.).

**Figure 3.**
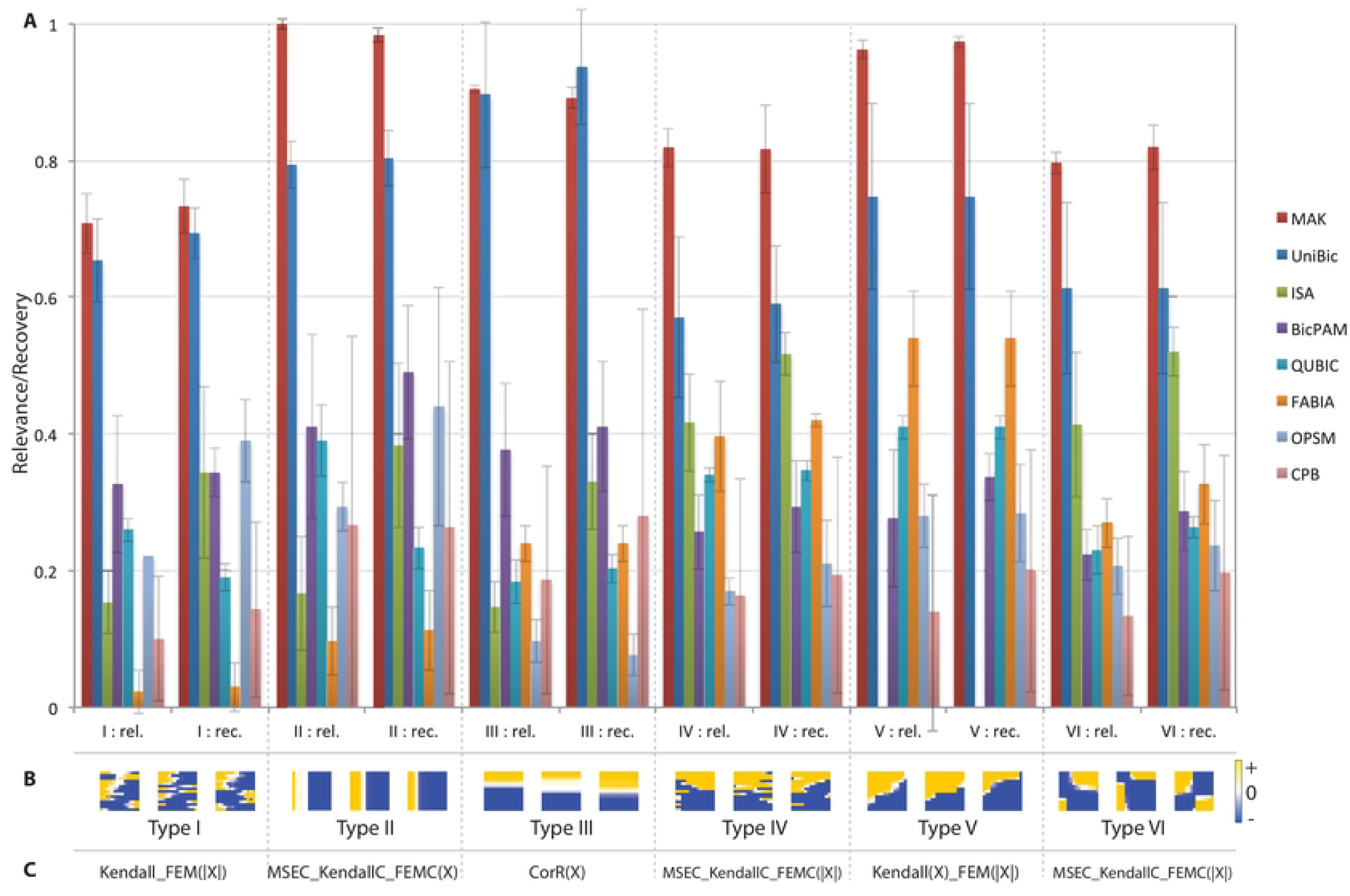
Evaluation of MAK on the UniBic simulated square bicluster data. We compared MAK to seven other biclustering methods based on the UniBic simulated data evaluation framework (52). This square bicluster simulated data spans six different bicluster pattern types and relevance and recovery values are shown for each biclustering method and error bars represent the standard deviation (A). EBIC and RecBic are not shown, see (44) and (45) for evaluation results. Methods are sorted by overall mean of relevance and recovery across all simulated data cases. The patterns of the simulated biclusters are show in in B and the applied MAK bicluster scoring criteria are shown in C. The absolute value of the input data is represented as |X|, and criteria ending in C or R are the column and row criterion version respectively, while no suffix indicates a criterion average of the column and row criteria.

### MAK *S. cerevisiae* gene expression biclustering results

A highlight of MAK parameters and command line arguments used to generate the yeast biclusters in this study is shown in Figure 4. We denote MAK results based on the specific bicluster coherence criterion used and on the data type layers which are being integrated, for example MAKcol(expr.) corresponds to column biclusters (“col”) from gene expression data (“expr.”). The MAK criteria used were determined from our high throughput simulated data evaluation (S1 Figure, Sup. Info.), chosen based on top performance and specificity towards a pattern type: KendallC_FEM for constant biclusters, MSEC_KendallC_FEM for column, and Kendall_FEM for row. We also applied a ‘checker’ pattern criterion by taking the absolute value of the input data for the column criterion MSEC_KendallC_FEMC. We focus on the combined MAK bicluster results for gene expression data across the three basic pattern types, column, constant, and row (denoted as MAKcol_const_row(expr.)), since they cover different types of bicluster patterns in a single result, and represent a nonredundant and maximum scoring subset of biclusters across pattern types. In the following sections we present evidence for the efficacy of MAK as well as for the selection of this MAK bicluster set as the exemplar MAK result.

**Figure 4.**
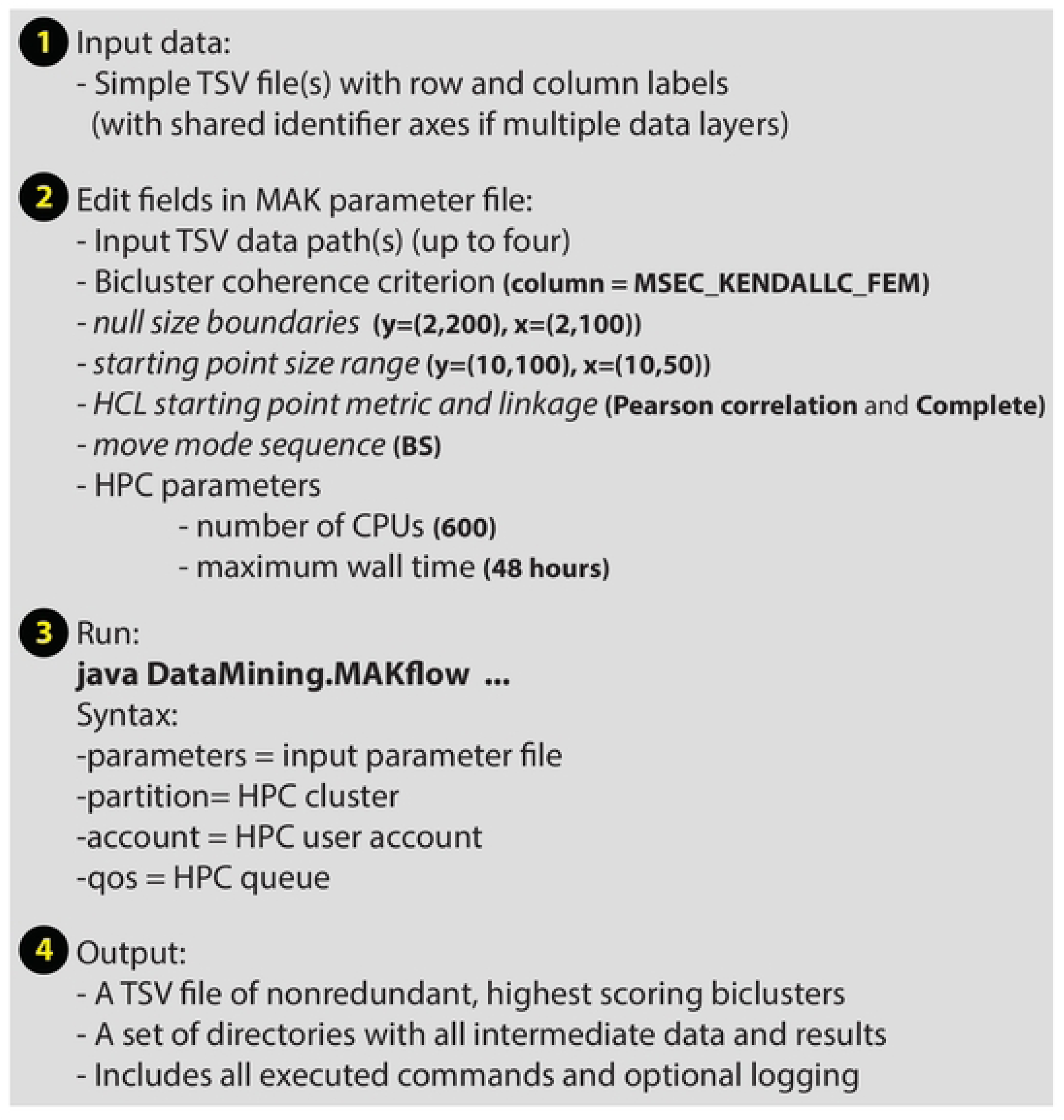
MAK parameters and command line arguments. An overview of the steps involved in running MAK biclustering. Core MAK parameters are shown along with example values used on the yeast data in this study. Finally the MAK pipeline is executed using a single command on a compute cluster.

### Evaluating biclustering methods with a data-driven score for real-world datasets without ground truth

The ultimate utility of a biclustering method is in its performance on real biological data. While for most biological datasets there is no established ground truth, properties of an ideal biclustering method can still be defined and assessed without knowing the underlying true patterns. Thus, an ideal biclustering algorithm should find the smallest set of the largest biclusters that covers the greatest portion of interesting values in the data, where the interesting values tend to be the most extreme. Ideal biclusters should also have high coherence, thus for a gene expression bicluster all data for genes and conditions belonging to the bicluster should conform well to the overall pattern. Biclustering methods that can find a balance of these important bicluster features are desirable and therefore we used these features to implement a bicluster set score not requiring ground truth (see Methods).

Table 1 shows biclustering methods sorted by their data-driven bicluster set score.

**Table 1.** Bicluster method comparison based on a bicluster set data-driven score without ground truth.

|  |  | Score | Correlation | Contrast | Condition coverage | Mean genes | Mean exps. | Number |
| --- | --- | --- | --- | --- | --- | --- | --- | --- |
| 1 | MAKcol_const_row(expr.) | 0.42 | 0.85 | 2.46 | 0.58 | 82.8 | 30.6 | 296 |
| 2 | MAKcol_const_row_checker(expr.) | 0.42 | 0.84 | 2.39 | 0.64 | 79.2 | 34.0 | 469 |
| 3 | MAKcol(expr.) Pearson-complete linkage | 0.42 | 0.87 | 2.51 | 0.51 | 86.3 | 23.2 | 246 |
| 4 | MAKchecker(expr.) | 0.41 | 0.83 | 2.24 | 0.36 | 72.0 | 41.0 | 181 |
| 5 | MAKconst(expr.) | 0.41 | 0.74 | 2.19 | 0.24 | 65.4 | 60.8 | 45 |
| 6 | MAKcol(expr.) Euclidean-Ward's | 0.41 | 0.87 | 2.40 | 0.76 | 49.6 | 14.4 | 1221 |
| 7 | MAKcol(expr.+TF) | 0.41 | 0.73 | 2.23 | 0.78 | 89.0 | 33.0 | 435 |
| 8 | MAKrow(expr.) | 0.40 | 0.83 | 2.26 | 0.14 | 89.9 | 85.0 | 15 |
| 9 | FABIA | 0.39 | 0.39 | 1.35 | 0.59 | 448.8 | 135.6 | 74 |
| 10 | EBIC | 0.39 | 0.87 | 2.12 | 0.69 | 383.7 | 11.5 | 100 |
| 11 | QUBIC2 | 0.39 | 0.82 | 2.99 | 0.21 | 84.7 | 15.7 | 100 |
| 12 | QUBIC | 0.39 | 0.80 | 3.34 | 0.05 | 45.8 | 34.9 | 10 |
| 13 | 2D-HCL Pearson-complete linkage | 0.39 | 0.67 | 1.69 | 0.53 | 60.8 | 31.9 | 150 |
| 14 | cMonkey2 | 0.38 | 0.47 | 0.88 | 1.00 | 19.9 | 326.7 | 616 |
| 15 | MAKcol(expr.+TF+PPI+ortho.) | 0.38 | 0.96 | 1.19 | 0.20 | 4.2 | 15.8 | 1199 |
| 16 | ISA | 0.37 | 0.55 | 1.76 | 0.40 | 114.4 | 35.2 | 72 |
| 17 | BicMix | 0.37 | 0.42 | 3.69 | 0.24 | 12.3 | 9.8 | 143 |
| 18 | cMonkey | 0.36 | 0.62 | 1.12 | 0.46 | 19.4 | 105.9 | 255 |
| 19 | COALESCE | 0.36 | 0.42 | 1.40 | 0.31 | 63.3 | 43.1 | 183 |
| 20 | 2D-HCL Euclidean-Ward's | 0.36 | 0.38 | 1.78 | 0.85 | 58.5 | 30.8 | 326 |
| 21 | RecBic | 0.36 | 0.44 | 1.16 | 0.97 | 1100.3 | 6.0 | 100 |
| 22 | RUniBic | 0.35 | 0.57 | 3.81 | 0.00 | 2.0 | 5.0 | 2 |

We use abbreviations to indicate the different integrated data types: ‘expr.’ for gene expression data, ‘TF’ for experimental TF-gene upstream region associations, ‘PPI’ for protein interactions, and ‘ortho.’ for yeast gene orthology conservation across related genomes. All MAK results spanning different bicluster coherence pattern criteria and data combinations, with one exception, ranked above all other biclustering methods. The exception were the MAKcol(expr.+TF+PPI+ortho.) results, which include three other data types in addition to gene expression. The gene expression contribution in these biclusters is expected to be diminished since gene expression and TF-gene associations account for only one half of the MAKcol(expr.+TF+PPI+ortho.) bicluster score components. Indeed, these MAK results had a high intra-bicluster correlation but 96% lower bicluster data contrast relative to average contrast across all other MAK results (contrast 1.19 vs. mean of other MAK results = 2.33). In addition, these MAK results resulted in smaller biclusters with on average 4 genes and 16 conditions. This is likely related to the imperfect agreement between the four data types used in this case, for example leading to biclusters, which exhibit weaker gene expression patterns but containing protein interactions or gene ortholog conservation. The MAK 2D-HCL starting points selected for more extreme differential expression (> 1.0 absolute mean gene expression) ranked below all MAK methods, with much lower mean bicluster correlation (22% less, 0.67 vs. mean of all other MAK results = 0.82) and contrast (31% lower, 1.69 vs. mean of all other MAK results = 2.33), indicating the improvement over starting points across all MAK results. The top non-MAK methods were FABIA, EBIC, QUBIC2, and QUBIC – with FABIA scoring higher on bicluster size properties features and EBIC, QUBIC2, and QUBIC scoring higher on correlation and contrast. For context, we show the top three biclusters from top biclustering methods ordered according to their MAK column bicluster coherence score (Figure 5).

**Figure 5.**
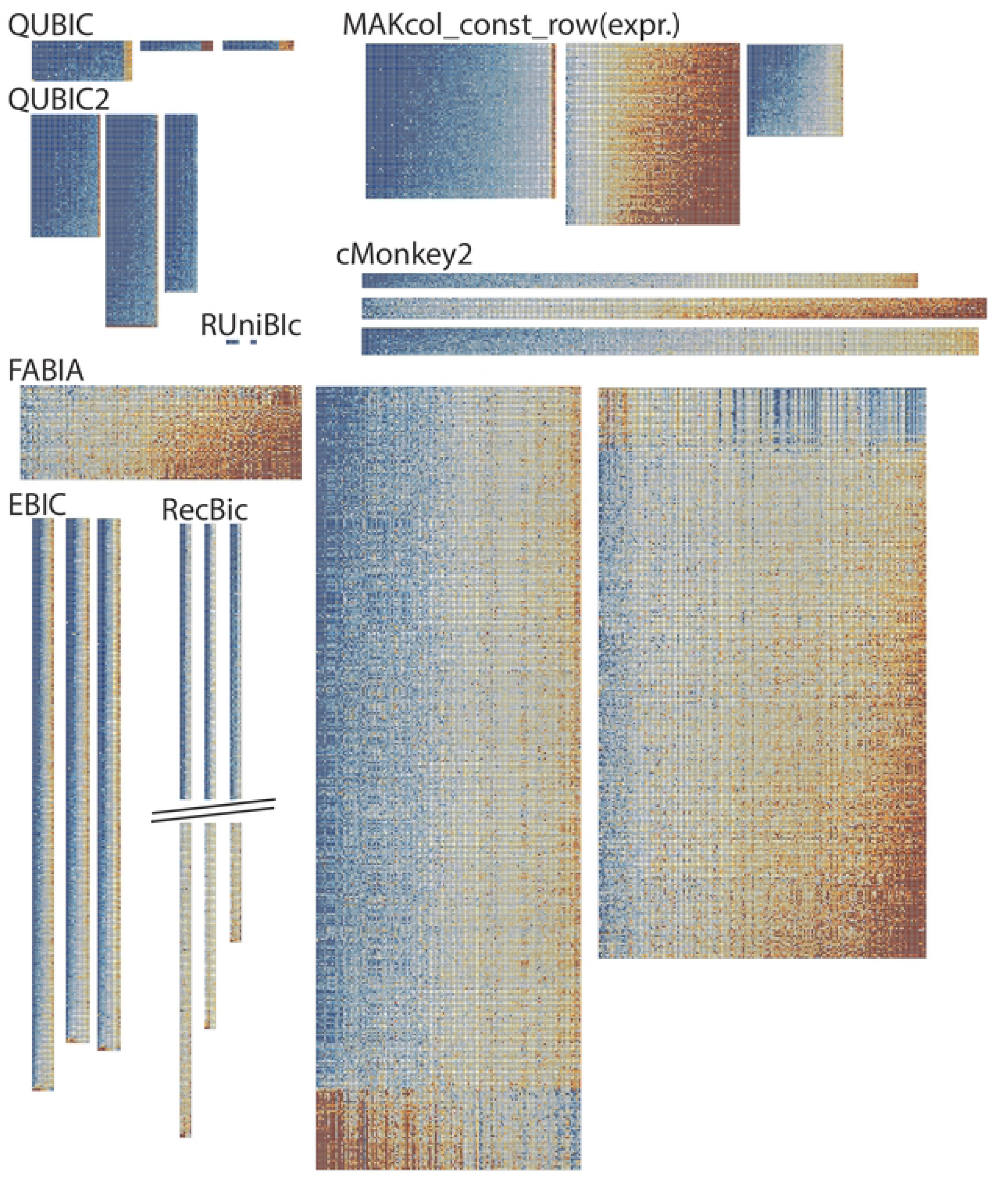
Example yeast biclusters from different biclustering methods. Shown are the top three yeast biclusters from each method, ordered based on their MAK column bicluster coherence score. Break marks for the RecBic biclusters indicate a portion of the biclusters that has been truncated for display purposes.

Methods are sorted by descending bicluster set data score, which is a Euclidean distance from a perfect score for the five bicluster set variables: mean correlation, mean contrast, mean gene coverage per number of biclusters, and the mean number of experiments. The values are colored column-wise using a red-blue (min.-max.) color scale. Column names with a gray background are components of the data-driven score. 2D-HCL results are filtered to include only biclusters with mean absolute differential expression > 1.0.

Additional bicluster features are shown in S1 Table, including the approximate runtime for each method on the yeast gene expression compendium. While MAK results in this study required the most compute time in CPU hours, the MAK pipeline results for a single pattern type like constant biclusters were only about 2-fold slower than other biclustering methods with a longer runtime that we evaluated (BicMix, cMonkey, cMonkey2). The MAK pipeline leverages high performance compute clusters and thus is inherently scalable by increasing the number of compute nodes and controlling the size and number of starting points, in addition to other algorithmic features (Sup. Info.).

In summary, the variables in the data driven bicluster set score allow to characterize biclustering methods solely based on the biclustering input data without the need to compare against ground truth. We do not include biological enrichment results in this score because for many datasets enrichment options are limited (e.g. only available for genes (i.e. GO terms)), the commonly reported enrichment percentages do not account for repeated enrichments for the same terms or term diversity or coverage, and relying solely on enrichment assumes that biological functions are well represented and covered in current classifications. In the following we focus on the top performing biclustering result representing all basic pattern types, MAKcol_const_row(expr.).

### MAK yeast biclusters have a wider distribution of number of conditions in biclusters

A major motivation for developing MAK was the presence of bicluster size biases in bicluster coherence criteria. MAK uses null distributions across bicluster sizes computed for each input dataset to explicitly adjust bicluster coherence criteria for size biases. We compared bicluster size distributions and statistics across biclustering methods on the yeast gene expression data (Figure 6, S2 Table). Different biclustering methods have different size characteristics. RecBic, EBIC, and FABIA where characterized by large numbers of genes, while cMonkey2 and FABIA were characterized by biclusters with the most conditions. FABIA, EBIC, and ISA also had the largest dispersion of number of genes in biclusters. The exemplar MAK results had the largest Inter Quartile Range (IQR) for conditions in biclusters, indicating the greatest statistical dispersion in the number of conditions. In addition the distribution of number of conditions in MAK biclusters was positively skewed, second after BicMix. Finally, of all methods, the mean ratio of number of genes to number of conditions for the exemplar MAK results was closest to the ratio present in the input dataset (39% of input ratio), just after QUBIC2 with a 42% ratio. The number of genes in biclusters should not vary dramatically based on input dataset size, especially when enough conditions have been sampled. This is because biological responses consist of finite signaling events deploying biological responses in the form of finite sets of genes and functions. However, the number of conditions in biclusters should depend on the number of conditions in the input data since the condition space is potentially large and partially degenerate with respect to biological activity and hence many conditions will map to the same or similar biological responses.

**Figure 6.**
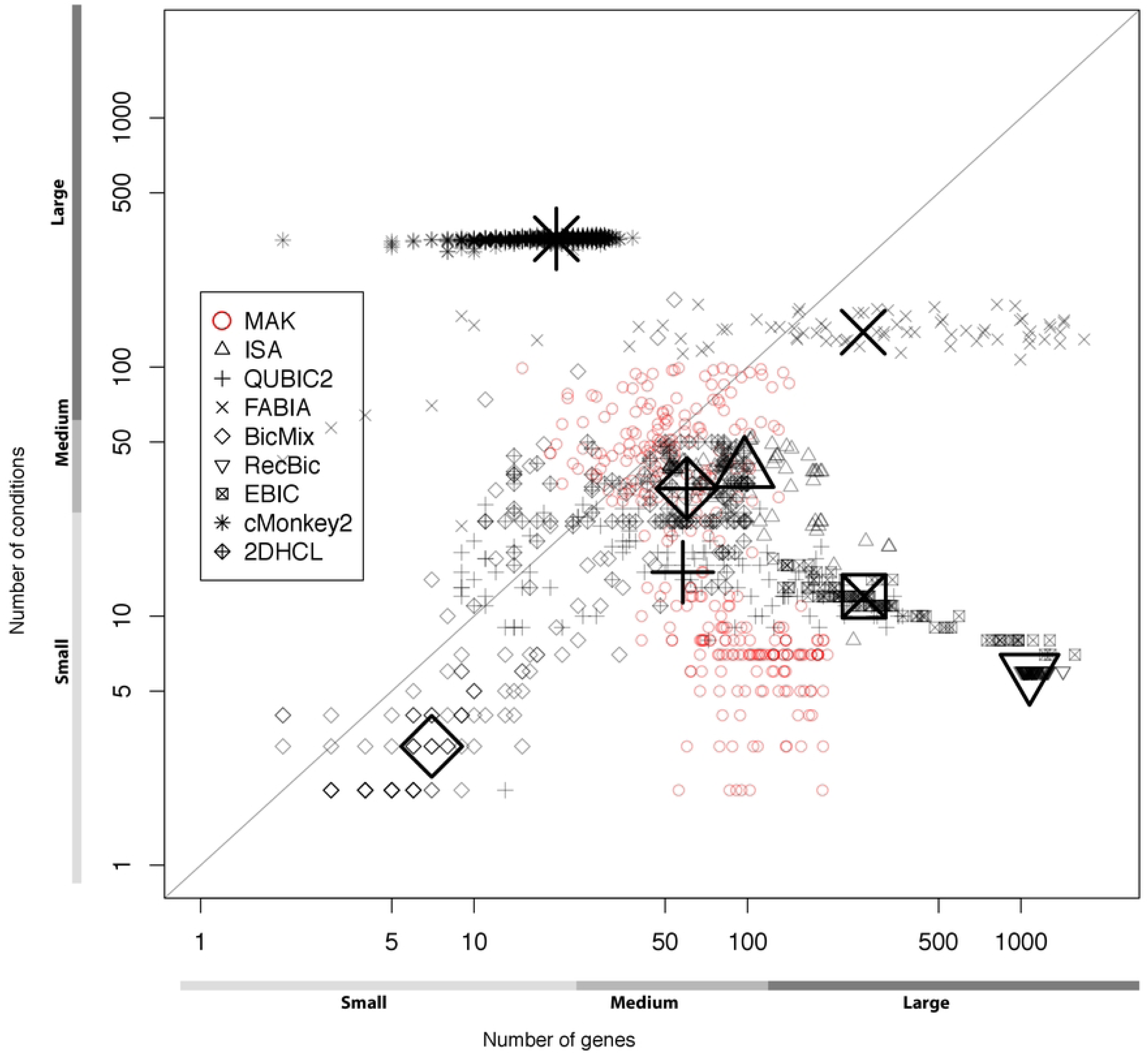
Yeast bicluster sizes across different biclustering methods. Biclusters from each method for the yeast gene expression data are plotted based on their gene and condition dimensions. Each method is represented by a unique glyph, and the median gene and condition bicluster dimensions are shown for each method with a single larger glyph. The x and y axis are on a log scale and the diagonal line represents x=y. The axis annotations show example regimes for different bicluster size ranges.

### Biological enrichment in yeast biclusters

We computed biological enrichments for all biclusters from the considered biclustering methods, using gene annotations with GO terms, TIGRFAM roles, KEGG pathways, and curated TF regulatory associations (S3 Table, Sup. Info.). MAKcol(expr.+TF+PPI+ortho.) had the highest coverage of GO terms with 43% of terms covered and the next top two methods were 2D-HCL Euclidean distance with Ward’s criterion (23%) and MAKcol(expr.) with Euclidean-Ward’s (23%). The top two methods for GO coverage also had the largest number of biclusters (1199 and 1221 respectively), thus for a fairer comparison we also compute per bicluster coverage (S3 Table). QUBIC, MAKrow(expr.), and FABIA had the highest per bicluster GO enrichment (4.4, 3.6, and 1.8 GO terms per bicluster, respectively). When considering the fraction of biclusters with enriched GO terms, three methods had GO enrichments for all biclusters (100%): EBIC, MAKrow, and QUBIC (S4 Table). All MAK gene expression results had a fraction of GO-enriched biclusters between 0.83 and 0.87, and only EBIC, QUBIC, RecBic, and FABIA had higher fractions of enriched biclusters. Since not all functions are currently known or well annotated, we may not expect perfect biological enrichment results for all biclusters representing true biological activity at this time and thus maximizing the fraction of enriched biclusters may introduce a bias against biclusters related to poorly characterized biological activity.

The top methods with the highest TF enrichment coverage were cMonkey (0.1), and 2D-HCL Euclidean-Ward’s and cMonkey2 (tied at 0.09). Out of all MAK results only the MAKcol(expr.+TF+PPI+ortho.) results with all data types had no TF enrichments, and all other MAK results showed TF enrichment coverage in the range 0.03-0.08. 2D-HCL with Euclidean distance and Ward’s criterion had 0.03 higher TF coverage than 2D-HCL with Pearson-Complete, but this difference was not observed in the corresponding MAK results, suggesting no effect of starting point method. When considering per bicluster TF coverage, QUBIC, MAKrow(expr.), and QUBIC2 showed the highest values ranging from 0.12 to 0.5.

The addition of TF 8mer binding data (MAKcol(expr.+TF) had no effect on the per bicluster TF coverage relative to gene expression data alone (MAKcol(expr.). MAK results with all four data types (MAKcol(expr.+TF+PPI+ortho.)) were the lowest in this regard with zero coverage, suggesting that inclusion of data beyond gene expression and TF associations can bias biclusters away from regulatory associations but also potentially related to our still incomplete understanding of cellular regulation.

The top three methods for KEGG pathway coverage were cMonkey2, 2D-HCL Euclidean-Ward’s, and cMonkey tied with MAKcol(expr.+TF+PPI+ortho.) (0.26, 0.18, and 0.17 respectively). Considering per bicluster pathway coverage, the top three methods were different: ISA, MAKrow(expr.), and MAKconst(expr.) (coverage 0.31 to 0.13). Finally, coverage of TIGRFAM roles was highest for MAKcol(expr.+TF+PPI+ortho.) tied with cMonkey, and with seven methods tied for second rank (FABIA, cMonkey2, COALESCE, three different MAK results, and the 2D-HCL Euclidean-Ward’s results). Per bicluster TIGRFAM role coverage was highest for MAKrow(expr.), QUBIC, and FABIA (0.13 to 0.07). Overall, the 2D-HCL Euclidean-Ward’s, cMonkey, and cMonkey2 ranked highest with coverage across the four functional assessments, while the MAKrow, FABIA, and MAKconst gene expression biclusters had the highest average ranking per bicluster confirming that MAK biclusters achieve more biological enrichments with fewer biclusters.

Considering only other methods, the exemplar MAK results MAKcol_const_row(expr.) ranked 4th, 3rd, 5th, and 2nd for unique GO, KEGG pathway, TF, and TIGRFAM role term enrichments, out of 11 top biclustering methods (excluding other MAK results and 2D-HCL). These MAK results were selected based on maximizing the bicluster set data-driven score, which did not include any features related to annotations with biological terms. Thus the exemplar MAK results represent the bicluster set with the highest data driven bicluster set score, close to the highest mean bicluster correlation (0.85 vs. EBIC = 0.87, Table 1), and performing better than all but three other biclustering methods on total coverage of biological terms from four different classifications enriched in biclusters (cMonkey (-0.33), cMonkey2 (-0.27), and only 0.02 less coverage than FABIA) (Figure 7), all of which exhibited considerably lower coherence. Since we lack a gold standard for biological enrichments there is a risk that gene sets from low quality biclusters can lead to spurious enrichment results with no associated biological activity. We attempt to address this problem by combining bicluster pattern quality analysis with enrichment results.

**Figure 7.**
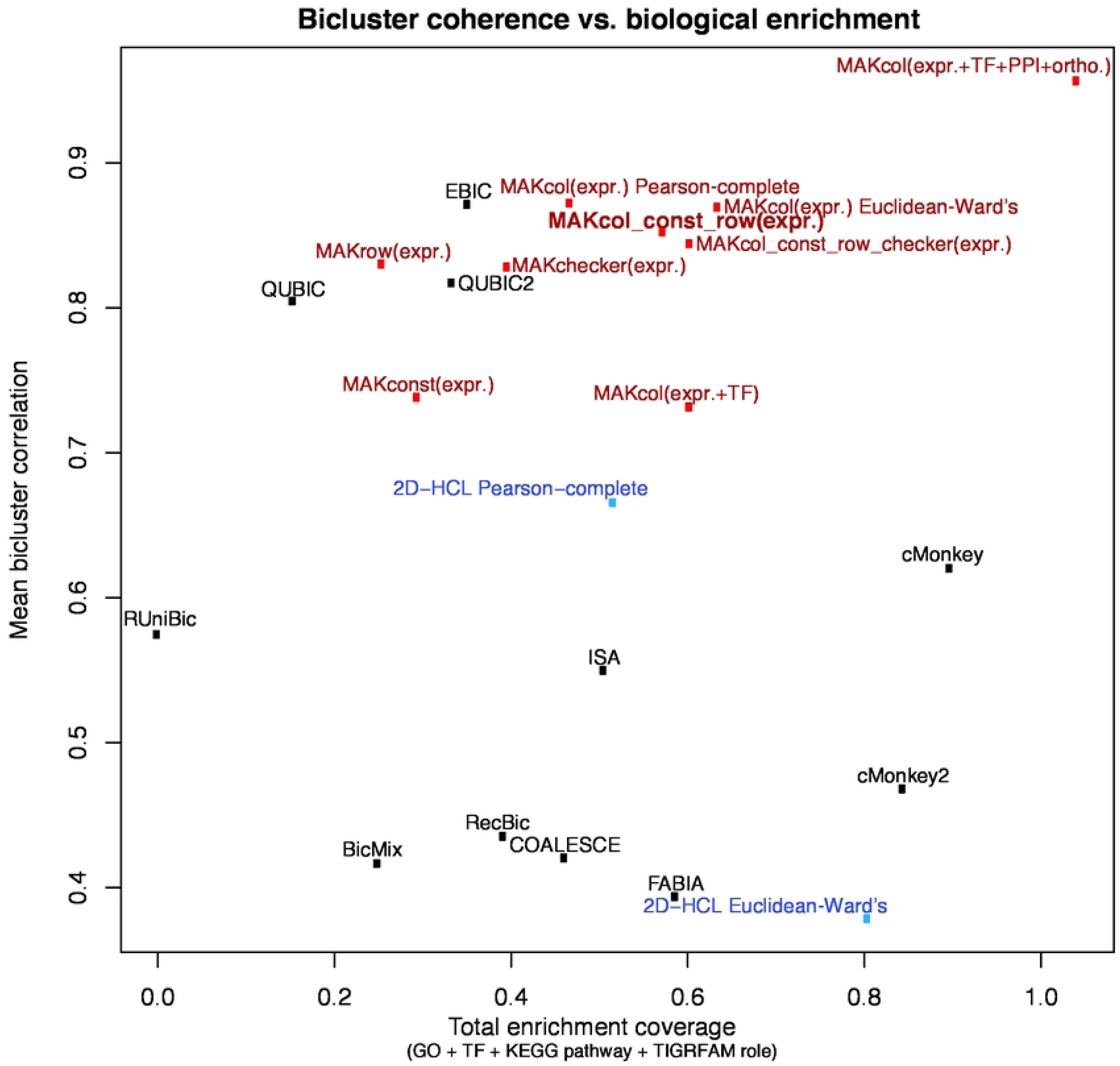
Relationship between bicluster coherence and biological enrichment. A scatterplot of the mean intra-bicluster correlation versus total biological classification coverage by statistical term enrichment in biclusters. Method and result labels are above or to the right of the corresponding data symbols. Bicluster sets are colored: MAK (red), MAK starting points (blue), and other methods (black). The exemplar MAKcol_const_row(expr.) results are labeled in bold. Enrichment was calculated for GO terms, YEASTRACT gene-TF associations, KEGG pathways, and TIGRFAM functional roles. 2D-HCL results are filtered to include only biclusters with mean absolute differential expression > 1.0.

### Yeast bicluster coherence and biological enrichment

For a broader assessment of biological relevance, we compared the total enrichment coverage of the terms from the four different biological reference data (GO terms, gene-TF associations, pathways, and TIGRFAM functional roles), to the intra-bicluster correlation from each method (Figure 7). The intra-bicluster correlation is reported as the maximum of the mean pairwise column, row, and total correlation, and serves as a measure of bicluster coherence. Total enrichment coverage was uncorrelated with bicluster coherence (R = - 0.014), across these 22 biclustering results. The range for the bicluster coherence was from 0.38 (2D-HCL Euclidean-Ward’s) to 0.96 (MAKcol(expr.+TF+PPI+ortho.))), while the total enrichment coverage statistic ranged from 0 (RUniBic) to 1.04 (MAKcol(expr.+TF+PPI+ortho.)). A few methods with very high enrichment coverage (> 0.8, i.e., 2D-HCL Euclidean-Ward’s, cMonkey, cMonkey2) had among the lowest coherence. The exception were the MAKcol(expr.+TF+PPI+ortho.) results, which had the highest overall coherence (0.96) and enrichment (1.04), indicating that it is possible to find bicluster sets with both high coherence and high biological enrichment. However, this MAK result utilized multiple high throughput experimental datasets, which may not always be available, and reported a large number of smaller biclusters with lower contrast.

Combining MAK results of different bicluster pattern types resulted in gains in both coherence and enrichment, including in the exemplar MAK results. To further summarize method performance we used the sum of the bicluster coherence and total enrichment coverage to rank biclustering methods. The top five methods were all MAK results with different bicluster pattern and data types. The exemplar MAK results ranked fourth, while the top method overall in this combined bicluster coherence and enrichment assessment was MAK using multiple data types, i.e. MAKcol(expr.+TF+PPI+ortho.). The pooled exemplar MAK results for three bicluster pattern types had greater coherence and biological enrichment coverage than any individual pattern type result, with the exception of a small decrease of coherence relative to MAKcol(expr.) results alone. Overall, these findings suggest that evaluating biclustering results solely based on statistical enrichments for biological terms, especially when using a single classification like GO, may not identify the best biclustering result. This analysis also demonstrated that combining high quality results for different bicluster pattern types and input data layers can lead to combined gains in both mean bicluster coherence and biological enrichment.

### GO term combinations enriched in yeast biclusters

To provide a more detailed picture of bicluster content across methods, we also calculated the distribution of enriched GO terms and GO term combinations (S4 Table). Two methods exhibited GO enrichment for all biclusters (QUBIC, EBIC) and the next top methods were MAKchecker(expr.) and RecBic, with 97% and 96% biclusters enriched for at least one GO term. Enrichment results can be convoluted by repeated enrichment for the same term or term combinations in a set of biclusters and thus higher coverage of GO ontology by enriched terms in biclusters is also desirable (assuming the biclusters also exhibit acceptable coherence). In this analysis every unique combination of enriched GO terms in a bicluster is treated distinctly. The total number of enriched terms per method reflects the coverage of GO terms by enrichment in biclusters, and the 2D-HCL Euclidean-Ward’s results contained 91 enriched GO terms with the next top methods being cMonkey (N=86) and FABIA (N=75). The 2D-HCL Euclidean-Ward’s results consist of somewhat smaller biclusters, relative to MAK results, as well as relatively high coverage of conditions in the data (85%, 3^rd^ highest, Table 1). The cMonkey results incorporated additional data types, which likely contributes to a greater coverage of GO terms via bicluster enrichment. FABIA biclusters tend to be larger and with above average condition coverage (Table 1).

The exemplar MAKcol_const_row(expr.) results ranked 6^th^ overall in total enrichment and were surpassed only by four different single bicluster type MAK results as well as EBIC. Since MAK results had the highest correlation and contrast of all methods, and the highest data driven scores (Table 1), there appears to be a biclustering tradeoff between bicluster coherence and bicluster biological enrichment, with MAK jointly maximizing these two key bicluster properties.

We compared the enriched GO term combinations across all methods using only gene expression data, by counting exact enriched term combinations (S3 Figure, S1 File). We excluded the 2D-HCL results and note that cMonkey used multiple data types (13) in addition to gene expression. This revealed that most term combinations are unique to a single method or MAK bicluster result (866/1137) and that no term combinations were shared between the 11 top biclustering methods. As expected, the MAKcol and MAKchecker results share more term combinations (9/1137) than MAKcol with MAKconst (1) and MAKrow (0). This is consistent with the bicluster pattern type model and MAK bicluster coherence criteria similarity, as the MAKchecker criterion is equivalent to the MAKcol criterion after taking the absolute value of the data. No GO term combinations were shared across the three individual MAK bicluster pattern types used for the exemplar results (column, constant, row). 46 term combinations were found by two or more non-MAK methods and 28 term combinations were shared between at least one MAK result and one of the other 11 methods. As expected, the greatest term combination overlap was for the MAK exemplar results and the individual component MAK criteria results MAKcol and MAKconst, with 177 and 12 shared term combinations respectively. The next highest overlaps were between the cMonkey2 and QUBIC2 results (11 term combinations), and then cMonkey2 and the exemplar MAK results as well as MAKcol (10 combinations).

Individual bicluster pattern MAK results did not share term combinations, with only MAKcol and MAKconst sharing a single term, indicating that individual MAK pattern types contribute orthogonal information. Overall, this combined enriched term analysis additionally validates the MAK results which have a term combination overlap similar to that among the set of 11 top methods. Furthermore, the nonredundancy of MAK results was confirmed at the level of putative labels for biological activities represented by the biclusters.

Hub effects can be observed in biological analysis, where a single or few biological entities or categories of biological processes dominate the results. Methods have been developed to remove data leading to hub effects via preprocessing (68, 69), however this can lead to a significant alteration of the original input data and should be undertaken cautiously. A hub effect has been often observed for the category of biological processes related to protein production, due to the importance of proteins in cellular processes as well as the volume of the cell required for protein production infrastructure. Results related to protein production often dominate biclustering studies however, at the expense of observing patterns for other biological processes or condition-dependent behavior of protein production related genes. One way to reveal hub effects in biclustering results is to measure the number of biclusters with functional annotations from non-hub categories. To this end, we curated a list of GO terms related to protein production (Sup. Info.) and summarized the bicluster biological enrichments after removing these terms (Table 2). The MAK exemplar results had the second most enriched GO term combinations distinct from protein production tied with FABIA at 48, with cMonkey2 having the most (N=66). These three methods had among the highest GO term coverage (FABIA (GO coverage = 0.19), cMonkey2 (0.18), and MAKcol_const_row(expr.) (0.18)), with only two other MAK results and the 2D-HCL Euclidean-Ward’s results being somewhat higher (coverage from 0.2 to 0.23) (S3 Table). Since MAK also exhibited the greatest mean bicluster coherence and contrast, considerably more than the other methods reporting many non-protein production term combinations, the MAK non-protein production biclusters are of higher quality and warrant further investigation. However, MAK also found the most unique term combinations associated with protein production itself (N=182), indicating greater detail and coverage of this core cellular activity. Overall, the MAK results had the greatest number of unique term combinations, nearly double the next method (230 vs. cMonkey2 = 128).

**Table 2.**
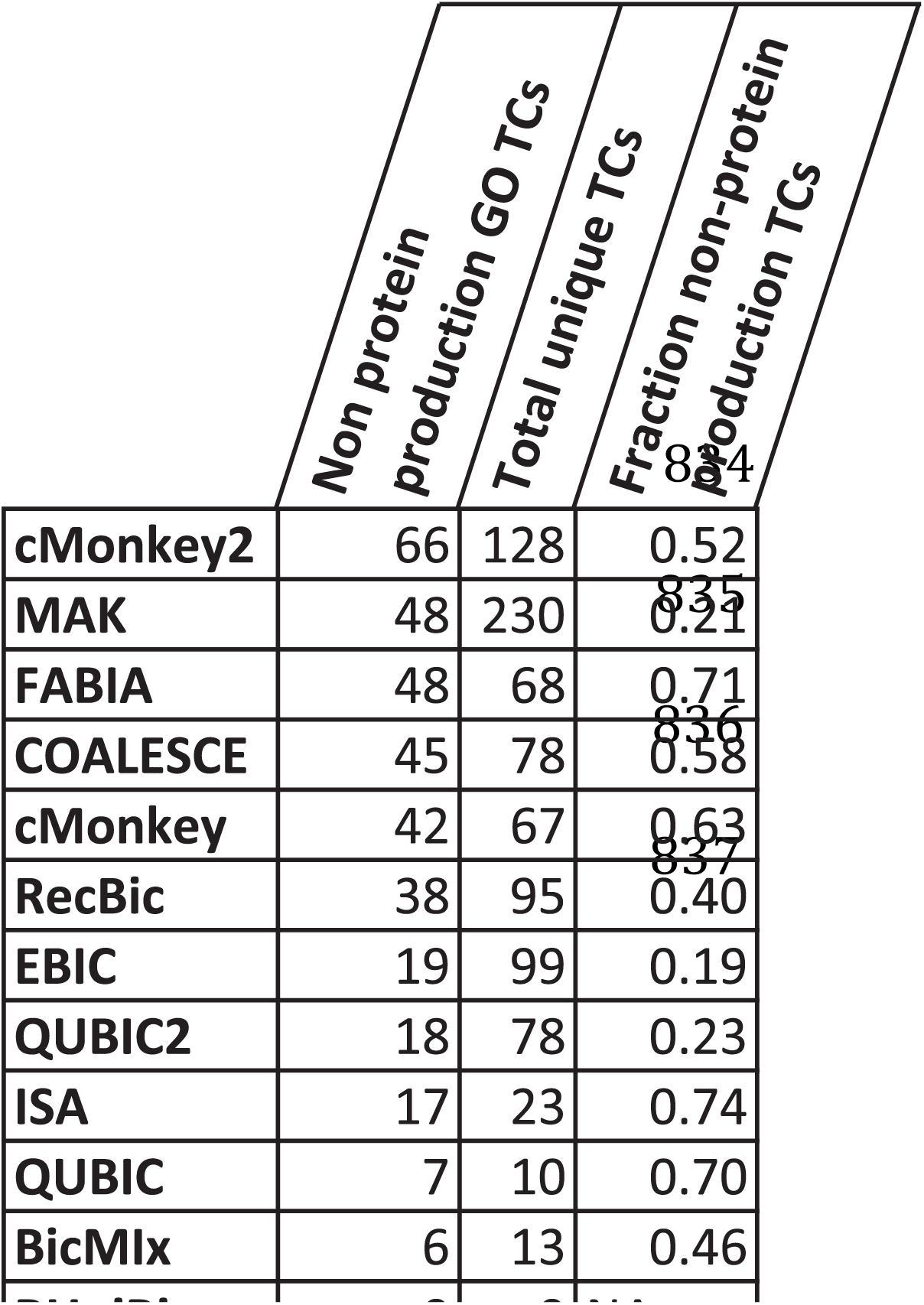
Enriched GO terms and combinations unrelated to protein production.

Most of these combinations are found in results from at least one other biclustering method, as the exemplar MAK results only represented 24 unique combinations, and these MAK results also had the fewest missing term combinations (907 missing compared to the union of term combinations from all methods, S4 Table). The large number of term combinations and high coverage of the union of all observed combinations, validated through results from other methods, and many biclusters found to be unrelated to protein production indicate greater overall diversity of biological functions represented in the MAK results.

A ranking of bicluster methods by their bicluster GO enrichments after removing GO terms related to protein production. MAK results are the exemplar MAKcol_const_row(expr.) bicluster set.

### Yeast bicluster overlap across and within biclustering methods

We compared biclustering results from MAK and other methods by measuring the fraction of biclusters exhibiting a non-trivial degree of overlap (Sup. Info., S4 Figure). We used a 0.25 overlap fraction threshold, corresponding to at least a quarter of bicluster area assuming equal bicluster sizes. The fraction overlap measure is asymmetric for two biclusters sets X and Y since the pairwise overlap data can be normalized by |X| or |Y|. The comparisons show that overall most methods share few similar biclusters and only the two small RUniBic biclusters are contained in many other results (overlap > 0.25 with 7/22 other methods). Hierarchical clustering of this overlap data reveals three main clusters on each axis, contained (x-axis) and containing (y-axis), and these sets of clusters exhibit some overlap. Both axes have a cluster of bicluster results which are both contained and containing each other, and are composed of mostly MAK results together with ISA, and one or two other methods. Biclusters from COALESCE, FABIA, QUBIC, QUBIC2, RecBic, cMonkey2 and MAKrow(expr.) and MAKcol(expr.+TF+PPI+ortho.) are not contained in any other method (except QUBIC2 in QUBIC and MAKrow(expr.) in the 2D-HCL Euclidean-Ward’s results). Similarly, BicMix, FABIA, MAKcol(expr.+TF), MAKcol_const_row(expr.), and 2D-HCL Euclidean-Ward’s are only contained, albeit partially, in FABIA biclusters. On the containing axis, cMonkey, QUBIC2, RecBic, RUniBic, and MAKcol(expr.+TF+PPI+ortho.) do not contain any other results, with only small overlap between cMonkey and RecBic with RUniBic. FABIA contains 13/22 methods or MAK results at maximum overlap > 0.25 with over half of their biclusters, consistent with the much larger size of FABIA biclusters (S2 Table). The FABIA results contain 63% of the MAK exemplar results with overlap > 0.25, suggesting that the considerably larger, less coherent and less contrasted FABIA biclusters may combine multiple smaller biclusters with higher individual coherence and contrast. We also note that the some FABIA biclusters have a checker appearance (Figure 5) where both a gene expression data pattern and its anti-pattern are included. This also helps explain the larger size of FABIA biclusters as well as the fact that they can contain smaller and more coherent biclusters of component types such as constant, column or row. However, checker biclusters can have better data qualities on this yeast data, for example the MAKchecker(expr.) biclusters were over 2-fold more coherent and with over 1.7-fold higher contrast compared to FABIA.

In addition, to understand the overlap of biclusters within each results, we analyzed the pairwise intra-bicluster fraction overlap for each method (S5 Figure), similarly focusing on more significant overlaps with overlap fraction > 0.25. Different methods have different approaches to biclustering and reporting biclustering results and for MAK we control bicluster overlap, and hence redundancy, by limiting overlap to 0.25 cosine similarity, meaning 25% overlap assuming equal bicluster sizes (cosine similarity is a symmetric overlap measure whereas overlap fraction is not). The bicluster overlaps in the MAK exemplar results were mostly restricted to the overlap fraction interval (0.1, 0.4) (median = 0.24). QUBIC (median = 0.15), QUBIC2 (0.38), FABIA (0.27), and RecBic (0.30) had overlaps in a similar range. The remaining methods had overlaps either in a higher range (> 0.4), indicating greater bicluster redundancy, or lower range (< 0.2) indicating marginal overlap. With these results we conclude that MAK biclusters exhibit plausible biological overlaps, similar to other top biclustering methods, and hence that MAK is able to recover overlapping biclusters in real data.

### Novel yeast biclusters discovered with MAK

The exemplar MAK results with the three bicluster pattern types using gene expression data alone identified many biclusters with little to no overlap with biclusters from other methods. To determine bicluster novelty, we used the asymmetric fraction bicluster overlap to measure the area of bicluster A covered by bicluster B and vice versa. The asymmetry accounts for differences in bicluster sizes, as the measures are normalized by the area of the left-hand bicluster in the comparison. Novelty required low (< 25%) to no coverage of a MAK bicluster across the 11 top biclustering methods (BicMix, COALESCE, cMonkey, cMonkey2, EBIC, FABIA, ISA, QUBIC, QUBIC2, RecBic, RUniBic). To identify the novel MAK bicluster subset, we required both coverage of the MAK bicluster and the MAK bicluster covering any bicluster from other methods to be less than 0.25 fraction overlap. Using the bicluster novelty criterion requiring novelty across 1655 biclusters from all 11 other biclustering methods, 21 of the 296 MAKcol_const_row(expr.) yeast biclusters were novel. The remaining 275 MAK biclusters were covered at least in 25% of their area by biclusters from other methods (S6 Figure). MAK biclusters were covered by other biclusters more than vice versa (0.33 vs. 0.22), except for 8 cases of full coverage (1.0) of other biclusters by MAK results. The highest coverage of a MAK bicluster by a bicluster from another method was only 0.74 (mean of the top 10 overlaps = 0.6), indicating partial overlap and differences in bicluster content. Thus the more coherent MAK results are only partially covered by biclusters from 11 other top methods, while MAK also returns additional novel biclusters with biological enrichments.

Data and biological enrichment features for the 21 novel MAK biclusters are shown in S2 File. These 21 novel biclusters were enriched for GO terms (10 biclusters), TIGRFAM roles (3), and pathways (1). In total 29 unique GO terms were enriched in this set of novel biclusters, including 9 terms not related to protein production (Sup. Info.): “actin filament”, “cellular respiration”, “generation of precursor metabolites and energy”, “mitochondrial respiratory chain complex II, succinate dehydrogenase complex (ubiquinone)”, “mitochondrial respiratory chain complex III”, “oligosaccharyltransferase complex”, “proteasome core complex, alpha-subunit complex”, “transmembrane transporter activity”, and “tubulin complex”. The three TIGRFAM role enrichments were “Protein synthesis” (N=2) and “Unknown function” (N=1). The single enriched KEGG pathway was “aerobic respiration, electron transport chain”. As the input gene expression dataset contained stress response experiments in various conditions, modulating protein synthesis and core energy metabolism is an expected key cellular activity and this is borne out by the biological enrichments observed in the 21 novel MAK biclusters.

### MAK for integration of multiple biological data types

For most non-model organisms, gene expression data is the most prevalent and sometimes the only available data. Hence reliable discovery of transcriptional modules solely based on this data type is important. However, integrating additional data types with gene expression can serve multiple purposes. One is to test hypothesis and generate models for specific biological mechanisms, for example gene co-expression patterns with underlying shared TF binding sites. Another is to help overcome noise in the primary dataset, here gene expression, by boosting bicluster scores based on coherence in other data types such as protein interactions and phylogenetic profiles. In the latter case, a gene with somewhat noisier expression data or affected by experimental bias but still weakly matching the overall bicluster pattern, can still be considered if it also co-occurs with these genes in the same genomes or if the corresponding protein interacts with other proteins in the bicluster.

Integration of four data types, namely gene expression, protein interaction, TF 8mer binding associations, and yeast pangenome orthologs, led to a decrease in the data driven bicluster set score (Table 1). In fact, these MAK results ranked as the lowest MAK results with the data driven bicluster set score. However, including the four data types led to some marked differences in bicluster properties: the highest mean bicluster correlation of all methods, lower coverage of conditions, low mean number of genes and conditions, and the 2^nd^ largest number of biclusters.

Considering the bicluster coherence and total biological enrichments (Figure 7), the multiple input data type MAK result outperformed all other biclustering methods as well as other MAK results. However, compared to integrating all four data types, biclustering gene expression data only with the TF 8mer binding associations led to a near doubling of the mean bicluster contrast relative to background (1.8-fold), as well as increases in condition coverage (3.9-fold), mean number of genes (21.2-fold), and mean number of conditions (2.1-fold). These features appear to involve a tradeoff between higher correlation, as the MAK results with all data types led to an increase in mean correlation relative to using expression and TF data alone (0.76-fold). The MAKcol (expr.+TF) results also had lower (1.7-fold) total biological enrichment relative to all four data types, indicating there is additional information in protein interactions and ortholog sequence conservation patterns. The latter is a data type, which can be universally generated from available genome annotations for many organisms.

The MAK results with only gene expression and 8mer binding data should correspond more closely to sets of TF targets under coordinated regulation. Indeed, the MAKcol(expr.+TF) results showed the highest enrichment for TF targets out of all methods and MAK results. A total of 42 biclusters from MAKcol(expr.+TF) were enriched for at least one TF, compared to 17 in the results with expression data alone and none in the MAK results with all four data types. This demonstrates that there is additional information in the experimental 8mer binding data that in turn leads to more enrichments for targets of those transcription factors, assessed with independently curated TF-gene target association data (see Methods). These MAK results were enriched for many more TFs, 42 vs. 11, than the cMonkey results (the only other method in our comparison, which included other data types, including sequence binding motifs). These results demonstrate how MAK can be used in computational experiments for targeted data and information integration studies.

Combining multiple data types corresponds to finding biclusters with a common gene expression response and with agreement in TF binding profiles for MAK(expr.+TF). For MAK(expr.+TF+PPI+feat.) the interpretation additionally involves similar phylogenetic distributions, and containing genes belonging to the same protein complexes. Phylogenetic profiles and interaction data have been suggested as synergistic for identifying transcriptional modules (13, 31), as are gene expression and TF binding. Biclusters with support in these additional data types can be interpreted as conserved protein interaction modules with co-regulatory associations or regulons with differential expression of protein complexes. Each dataset is incomplete to different degrees and in different ways, and furthermore the agreement between data types is limited by relationships defined by biological mechanisms.

## Discussion

The MAK bicluster discovery strategy includes innovations such as data adaptive null distributions to account for bicluster criteria size biases, comprehensive coverage of the input data with informative starting points, performing many parallel searches with HPC, as well as nonredundancy and merging procedures to derive information from overlapping biclusters, including for bicluster sets representing different bicluster pattern types and input data combinations. These functionalities are contained in the MAK framework and include bicluster scoring criteria, methods to compute null distributions, the MAK algorithm itself, a simulated data test-bed for high throughput bicluster coherence criteria assessment, components to deploy MAK HPC bicluster discovery strategies, and post-analysis tools. The framework provides a platform to perform cycles of biclustering and assessment, allowing to optimize criteria and search strategies for individual datasets given that no single biclustering method or criteria are expected to sustain performance across different data sets and data types (40).

While MAK with gene expression data alone jointly maximized bicluster correlation and total enrichment compared to other methods, we observed the best performance overall with MAK and the combination of gene expression with three other data types.

Generation of each dataset incurs a significant experimental cost and thus these are unlikely to be available except for model organisms. We observed a performance discrepancy between MAK results with gene expression data alone, gene expression data with experimental TF binding data, and MAK with four different data types. In particular, the most coherent gene expression biclusters mostly do not correspond to known TF-gene regulatory associations, as evidenced by the relatively low enrichment for TFs across the methods considered here. EBIC results were the exception, as 95% of biclusters were enriched but only repeatedly for the same five TFs. There is value in both identifying the most coherent biclusters in a single data layer as well as across data layers, though these are different formulations of the biclustering problem and can give different results. These results can also have different applications, for example gene expression data alone can identify sets of genes responding in a coordinated fashion but unlinked from current knowledge about transcriptional regulation. It remains to be seen whether improved knowledge about regulatory processes and condition-dependent cellular behavior can better explain the observed coherent gene expression data patterns.

The true number of biclusters in a biological dataset is generally unknown, therefore biclustering methods apply thresholds or treat this variable as an input parameter (13). Some methods may be better at finding small very coherent biclusters, while others may excel at finding larger but less coherent ones – these differences may affect how a larger pattern can be split into multiple patterns with higher coherence. MAK uses a novel algorithmic approach and estimates the true number of biclusters statistically with a large number of potentially overlapping biclusters, which are selected by minimizing overlap and maximizing the score. In reality, the true number of biclusters in a dataset is determined by multiple factors including the number and type of conditions or features, the underlying structure of the data (e.g., the transcriptional regulatory network for gene expression data), as well as algorithmic features affecting the performance and nature of bicluster recovery. The latter may lead to missing patterns, false patterns, and merging or splitting of biclusters belonging to larger and potentially less coherent patterns.

Since the number of biclusters is dependent on the input data, the bicluster pattern model, and the algorithm implementation, and because in general true patterns are unknown, we have to be able to compare biclustering results in a bicluster number and size-independent manner. The asymmetric bicluster set overlap analysis is one such approach, which can reveal details about the correspondence and relationships between different biclustering results. We observe that biclustering performance can degrade and the nature of biclusters can change with increasing types and numbers of data layers, resulting in different sizes, numbers, coherence, and contrast of biclusters. The growing number of biological datasets and data types is leading to a combinatorial explosion of data combinations as well as potential hypotheses, thus comparing similar or overlapping biclusters of different natures and sizes will remain an important problem.

Due to its data-adaptive aspects, MAK is less likely to require data preprocessing, but we assume that the rows and columns are normalized appropriately for a given data type and the row or column means are approximately zero, as in this study. Although methods exist to adjust data to account for batch effects (70), there have been no assessments of batch effects on biclustering nor of effects from other data normalizations. Future studies should address whether pre-processing can mitigate potential batch effects and ensure that for example different genes are comparable across different conditions.

However, since MAK relies on a combination of statistical criteria and employs a data adaptive null distribution, both normalization and batch effects are expected to be mitigated since each the bicluster moves, the addition or removal of each column or row, are assessed independently.

Biological enrichment results play an important role in assessing results on real-world data without ground truth, thus it is worth exploring improved approaches to performance enrichment and term analysis. For example, semantic similarity measures could provide a more refined similarity measure to compare enrichment results between biclusters as well as bicluster sets from different methods, by measuring similarity between sets of enriched terms in biclusters. This could also be used to unify enrichment results across different biological reference data, e.g. gene functions from GO and pathways from KEGG. We showed that bicluster GO term enrichment combinations are a more sensitive measure of bicluster functional similarity than individual terms or bicluster enrichment percentages. Combined with similarity measures, which can measure semantic overlap, such approaches can advance the interpretation biclustering results and method evaluations. Biological enrichment analysis can be sensitive to the exact content of biclusters and moreover can be affected by the enrichment test methodology and applied thresholds, for example (71). Therefore, GO enrichment results alone do not suffice to compare and evaluate biclustering methods and in fact may be misleading if not placed into a bicluster coherence context. More advanced statistical tests can be performed on enrichment results such as computing enrichment for randomly sampled matrices of the same size (e.g. (72)). Future work will address detailed comparisons of bicluster overlaps and their features, such as enrichment specifically for overlapping bicluster intersection and the remaining ‘overhang’ data from each bicluster in the pair. Finally, recent biclustering evaluations only consider bicluster or cluster gene content (52, 72) because it is difficult to evaluate biclustering results using metadata for column features like samples or conditions, due to a lack of standardization and supporting classifications. However, including column features in method evaluations is important, can support metadata enrichment analysis, and will help distinguish biclustering method performance from clustering methods.

More realistic simulated datasets for biclustering evaluations are also desirable, perhaps inspired by well-understood experimental data compendia. Currently there is a gap between results obtained with simulated data and performance observed on real data – in summary, simulated data performance is not very predictive of real-world data performance. Unfortunately this gap is preventing direct application of top methods on simulated data to real world examples. One difficult unaddressed problem is the generation of simulated data together with biological annotations with reference data like ontologies, which will require additional modeling. This would allow measuring method performance more closely to how we expect to measure it on real datasets without ground truth.

Otherwise, it will remain difficult to validate biclustering results simply based on isolated biological enrichments, although computing enrichment against multiple independent biological classifications should help alleviate this problem. In addition, nuances involving the exact statistical enrichment methods applied, the effect of applied significance thresholds, as well as issues with under- or over-annotation of the input data, all can contribute to observed differences in biclustering performance and interpretation.

Standardization of annotation and enrichment methods should contribute to resolving some of these problems. At the moment, researchers are mostly unaware of data and method biases and how these affect method performance such as biclustering. More studies are necessary for comparing biclustering results across different datasets to gain an understanding of factors affecting performance.

We envision further improvements relative to the current MAK concept and implementation. In MAK, the random seed can affect the starting points and the content of the batch moves but it can also be used to randomize the move type sequence, or to add a Metropolis move criterion. Other algorithm modifications are possible, for example allowing to decrease the number of sequential rounds of bicluster searches through a larger number of parallel but divergent trajectories. Here we considered a simpler approach, which uses a pseudo-random walk to bias towards larger biclusters and gene moves. Compared to a random walk, this choice of a move type sequence also renders the trajectories from the same and different rounds more comparable. Other types of improvements include pre-computed null distributions for popular datasets. An analysis of null distribution shapes along with exploring different parameterizations across multiple data sets and data types could reveal more general principles and allow these distributions to be computed analytically or to be learned more efficiently. In addition, various aspects of the source code can undergo optimization, both at the criteria, the algorithm, and the bicluster discovery pipeline levels. These improvements are well supported by the current MAK framework, which implements a workflow that can be restarted at any step or reused and forked in future analysis, and moreover can save all intermediate results, executed commands, and along with logging. The MAK implementation is a tested and hardened computational pipeline and only requires access to a modest compute cluster or even a single CPU for targeted searches using pre-computed nulls.

Given the number of biclustering methods, the number of demonstrated applications may still be underwhelming. A few trends should increase the relevance of biclustering. First, datasets are increasingly generated in more standardized ways and with better experimental techniques. Second, the importance of metadata and standardized labeling is gaining wider acceptance. Finally, knowledge contained in reference data such as ontologies is increasing, and the methods used to apply this knowledge to generate e.g. gene annotations, are also improving. There are many biclustering methods, however with somewhat greater resource requirements, MAK can provide the highest quality biclusters also reflecting desirable biological properties, including term enrichments, a wider dynamic range for the number of conditions in biclusters, as well as overlapping patterns across multiple bicluster pattern types. Surprisingly, on a medium yeast gene expression dataset, MAK was able to find 21 biclusters not found among 1655 biclusters from 11 top methods. As each biological unit of knowledge has potential value, MAK can provide comprehensive and high quality biclustering results above and beyond combining results from many state of the art methods.

The advances represented by MAK and the accompanying method evaluation should contribute to wider adoption of biclustering and biclustering results. These advances can also help overcome existing machine learning challenges in biology involving dimensionality reduction and pattern identification. Future work will address formal evaluations of multiple data type integration and as well as further improvements in scaling and efficiency.

## Methods

### Bicluster scoring criteria

We distinguish a block, which is any submatrix of the larger dataset, from a bicluster, which is a special case of a block also exhibiting a significant pattern. The details of the pattern are determined by the data type in question and the applied bicluster coherence criterion and algorithm. In this study we focus on constant, constant column, constant row, and the composite checker bicluster pattern models.

### Gene expression criterion: MSE

Criteria based on the MSE provide a measure reflecting bicluster consistency where consistency is defined as low deviation from the mean. They can measure full consistency (MSE) or row or column consistency (i.e. MSER, MSEC). Unmodified criteria from the MSE class will tend towards more favorable values for smaller blocks, which we address by computing scores based on a null model (see below). We compute the row or column MSE as a fraction of the total MSE, for example for MSEC:

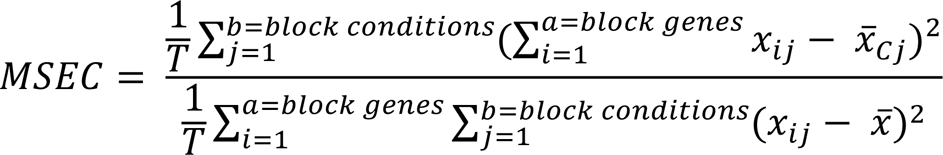

Where T is the total area of the block, 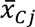 is the mean of the column j in the block, and 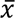 is the mean of the block. MAK bicluster criteria scores range from 0 to 1, where 1 is a perfect score. Thus the MSE criteria were transformed to the (0, 1) interval, where 1 is no deviation from the mean, by

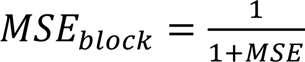

### Gene expression criterion: Kendall’s W

Criteria based on Kendall’s W non-parametric statistic provide a measure reflecting bicluster consistency based on agreement between voters, where voters are an example of more general decision-making. Voters are represented on one axis of the data and correspond to genes or conditions in gene expression data. We use the coefficient of inter-voter reliability as the test statistic. Classification decisions for bicluster membership are determined based on gene or condition expression profiles, for row and column bicluster scoring criteria, respectively. For example, the column version of this criterion is:

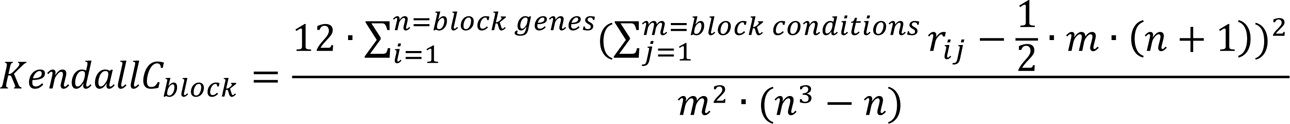

In general, *n* is the number of objects, *m* is the number of voters, and *r_ij_* is the rank of object *i* by voter *j*. For the gene expression data column criterion, the raters are conditions and the objects are genes.

### Gene expression criterion: fixed effects model (FEM)

The fixed effects model bicluster criterion was developed to provide a contrast metric that measures how well the row or column profiles within the current block differentiate themselves from the rows or columns outside the block. Specifically it compares the row or column means within the block to the overall mean outside the block using a regression model format in which the outcome variable is the profile of expression values and the covariates are indicators for row or column membership. For example, if one is interested in a block where each column has similar means, the set of full data profiles for these columns (including all rows), can be formed into a longform vector (*Y*). This vector can then be regressed on a set of indicator variables (*X*). Each indicator variable within X represents a specific column in the original block and will have a value 1 indicating membership of that column and zero otherwise. The criterion value is the coefficient of determination (R^2^) resulting from the regression fit given by the equation:

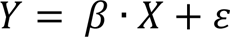

This bicluster criterion measures how well the difference in condition means determined by the block boundaries (i.e., mean over genes inside versus outside the block) explains the variation in gene profiles inside and outside the block over the conditions. This can also be determined with respect to the other axis, considering a set of condition profiles over a set of genes were the resulting metric represents how well the difference in row means determined by the block boundaries (i.e. mean over conditions inside versus outside the block) explains the variation in condition profiles inside and outside the block over the genes. The criterion value is given by:

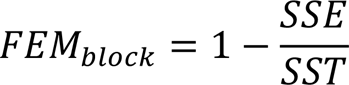

Where the sum of squared errors (SSE) is the sum of squared differences of row or column values and the row or column means in the block and outside the block, for the selected columns and rows respectively. The total sum of squares (SST) is the sum of squared differences of deviations of values in the block relative to the mean of the block. For, example FEMC is given by:

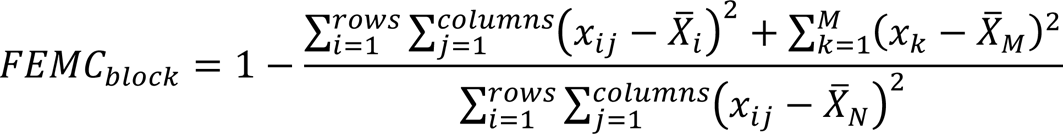

Where x_ij_ are values in the block, x_k_ are values outside the block and M is the total number of values outside the block, and x_k_ are values in the block and N is the total number of values inside the block. In most of our simulated data evaluation scenarios we found that subtracting row means showed improvement for column pattern recovery and vice versa for row patterns. For column biclusters the FEM criterion considers the errors for genes in and out of the bicluster restricted to the conditions in the bicluster and vice versa for row biclusters. We compute the FEM criterion for the absolute value of the data, after row or column missing data imputation (for row and column criteria, respectively).

### Graph density criterion

For binary interaction data, i.e. protein-protein interactions, we use a proportion of interaction criterion related to graph density. This value is calculated as

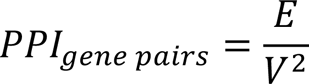

where E is the number of edges and V the number of vertices. This formulation allows for non-symmetric interaction data, which is expected in protein pull-down experiments since each bait protein needs to be tagged independently. This measure also allows for self interactions.

### Hamming distance feature criterion

We used the mean pairwise Hamming distance as a criterion for coherence of a set of genes (or condition) in a binary feature dataset, such as phylogenetic profiles. In this study the binary feature data corresponded to gene phylogenetic profiles across 79 fungal genomes. This criterion was transformed to the (0, 1) interval, where 1 is perfect agreement, as follows:

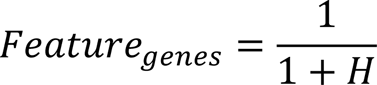

### TF-gene association rank criterion

For the experimental transcription factor gene target rank data we use the ranked 8mer microarray intensity profiles from (73). Each bicluster is evaluated based on the profile of mean TF ranks for the genes in the bicluster. The criterion value is the median of the minimum TF mean ranks for the set of genes in the bicluster, normalized by the maximum mean rank. Two minimum TF ranks are considered by default.

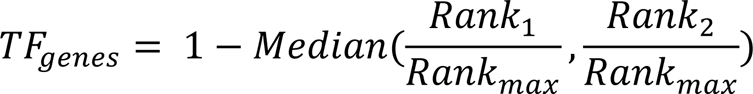

### Maximizing the fraction of row or column mean sign agreement for constant biclusters

For criteria targeting constant biclusters we implemented an additional criterion modification that measures the fraction of columns or rows or columns and rows, which agree on their sign based on the column or row means, respectively. The column, row, or column and row sign agreement was applied to the corresponding column, row, or total constant bicluster pattern type criteria. The sign agreement fraction was used to scale the raw criterion value and null distribution parameterization was performed on this scaled value.

### Null models for bicluster scoring criteria

Given that many comparisons occur at the extremes of the criteria value distribution, it is not feasible to use empirical null models for our criteria. Instead we chose to approximate the null distributions using a Cauchy distribution because of its ability to model heavy tails, allowing comparisons further out in the tails of the distribution. We used the sample median and 0.5 of the interquartile range (IQR) as parameters of the Cauchy distribution. Though preliminary assessments using density plots did indicate adequate fits, the Cauchy distribution is theoretically not an ideal distribution for many of the bicluster scoring criteria and future MAK versions will consider alternative distributions, specifically evaluated for each criterion.

As a way to decrease the computational demand for the null model generation, blocks with valid dimensions were sampled in five dimension bins for genes and conditions. Sampling density was decreased linearly from the first to last increment, that is for each increment *i* in 1…5 the increment number determined the sampling density within that increment such that in the first bin (2, 20) sampling was performed for each gene size, in the next bin (20, 40) every second gene size and so on. We generated 100 sets of 50 samples for each bicluster criterion and each sampled bicluster size. Next, the mean of the median and 0.5 IQR of bicluster criteria values over the total 5000 samples was computed for each sampled bicluster size. The mean summary statistics for all valid bicluster sizes in the specified range were obtained by fitting a thin plate spline to the sampled grid data. We used the linear model resulting from the thin plate spline to predict the summary statistic values on the expanded grid. For gene-only criteria, including PPI, TF, and gene features, only the gene samples were used from all of the blocks samples for the gene-condition criteria.

Based on the sampled bicluster criteria values we generated a dataset specific null model for each combination of a bicluster criterion and an allowed bicluster size range. Using the median and 0.5 IQR values given by the predicted values from thin plate spline linear model we generate null models based on the Cauchy distribution. With these null models we obtain scores for bicluster criteria values. The scores place disparate criteria, such as R^2^ values and MSE measures, on a common scale, thus allowing rigorous comparison and combine different criteria and data types. In addition, the null model allows to overcome size biases of bicluster criteria, such as the tendency of the MSE to achieve minimum values through minimizing the size of the bicluster.

### Algorithm

The MAK algorithm performs moves in bicluster space by accepting or rejecting valid moves relative to the current block (Figure 2.A). Blocks already visited in the trajectory are no longer considered valid moves. If from the candidate block list any block has a criterion score greater than the current block, this maximum block becomes the new current block. If there is no block with a criterion score greater than the current block, then the algorithm chooses a remaining move type and move direction. The selected move type determines the list of candidate moves, which are available starting from the current block and satisfy the missing data requirements. We use a default missing data tolerance of ***M_g_***=20% for genes, ***M_e_***=20% for conditions, and ***M_t_***=10% for the total block. A move type is skipped if the null model bounds are reached for block size, if the possible moves do not satisfy missing data thresholds, or if no moves of that type can improve the score of the current block. If there are no more remaining move types possible then the search completes and reports a final block.

At the core of the MAK algorithm are two types of sequences, which specify how the search in the bicluster space will be performed. The first sequence consists of an ordered series of move modes that the algorithm will attempt during the search. The default settings are batch moves followed by single moves. MAK makes available additional algorithm move modes. A ‘random’ mode consists of a one-time addition of random genes and/or conditions up to ***F_flood_*** percent of the size of the current block. In ‘plateau’ mode suboptimal moves, ones whose criteria values are lower than the current block, are accepted based on a specified noise threshold. Finally, there is a refinement mode during which the algorithm repeatedly performs single moves using modified convergence rules, until absolute convergence, and scoring larger blocks using the null model parameter distribution edge values. The ‘single’, ‘batch’, ‘random’, ‘plateau’, and ‘refinement’ move modes can be combined in any sequence and order and in this work we evaluate use of ‘single’ and ‘batch’ moves. For each move mode there is the possibility of using a pre- or full bicluster criterion. The algorithm stops when it reaches convergence, i.e. when all move types in the move type sequence have been tried for all move modes in the move mode sequence. Convergence is relative or absolute, depending on the move mode and parameters. The move types are the other type of sequence in the MAK algorithm and correspond to moves which are data type-dependent. For gene expression data these moves correspond to gene addition, condition addition, gene subtraction, and condition subtraction. These same move types are used in both the batch and single move modes, with the former moving based on multiple and the latter on single genes and conditions.

The single move mode can be computationally demanding for exploration of large data spaces hence we additionally designed a batch move consisting of addition or subtraction of multiple genes or conditions. Batch moves are allowed if the current block size permits adding or deleting a minimum number of genes or conditions (default = 3) and when the move does not alter more than a fraction of the bicluster data (batch move proportion parameter of 20% for genes or conditions by default). The candidate blocks corresponding to valid batch moves are clusters of genes or conditions obtained from hierarchical clustering, with the user-defined clustering parameters mirroring those used to generate the initial starting points (e.g., by default HCL with (1 – ((Pearson correlation+1)/2)) and complete linkage). The clustering is applied to all genes or conditions not in the current block (addition) or all of the genes and conditions in the current block (subtraction), restricted to the conditions or genes in the current block respectively. Missing values are imputed for each block with the mean for the row or column of the block. Distinct clusters, including singletons, of genes and conditions used to generate the candidate move list are obtained by cutting the clustering dendrogram based on a cluster size heuristic. The heuristic preferentially increases the maximum size for blocks with more genes or conditions, with 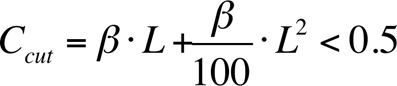 where β is the batch move proportion parameter and *L* is the number of genes or conditions in the block. Smaller values of β may help prevent trajectory drift, where the endpoint is completely unrelated to the starting point. It may also inhibit attraction to large biclusters with high criteria values in which case adding a small fraction of the attractor bicluster to the current block can significantly increase the criteria values and potentially mask the original pattern, especially for weak starting points. The choice of the height at which to cut the clustering dendrogram is computed using a bisectional search starting at the midpoint height. As with any candidate move, batch moves not satisfying the maximum allowed missing data thresholds are discarded.

The algorithm can use any subsets of gene and condition indices from the dataset as a starting point. Genes and conditions falling above missing data thresholds are trimmed from the starting point before any moves are considered. The algorithm can use a prescribed move sequence or can stochastically choose a move type based on specified initial move and subsequent move probabilities or use the combined approach where a move sequence is stochastically chosen from a set of prescribed sequences as we do in this study. The MAK move type probabilities are reset every time a move mode transition occurs, e.g., when ending batch moves and starting single moves or when restarting the single move mode during refinement. To direct the MAK random walk through bicluster space, the search trajectory is composed of a series of moves originating from repeated attempts to perform fixed sequences of move types. The first move sequence in a trajectory always starts with gene addition but subsequent move sequences first randomly choose to delete or add a gene. Every gene move is followed by the corresponding condition move and moves continues until all available moves in the move sequence have been attempted. Additionally heuristics can be used such as probabilistically balancing gene and condition moves, by setting gene versus condition move probabilities based on the proportion of the number of genes relative to the sum of the number of genes and conditions.

The full bicluster criterion is a weighted sum of a combination of bicluster criteria scores given by the null models, including gene-by-condition, protein-by-protein, gene-by-TF, and gene-by-feature. For results in this work corresponding to three different combinations of data types, the criteria are weighted as follows (Equation 1):

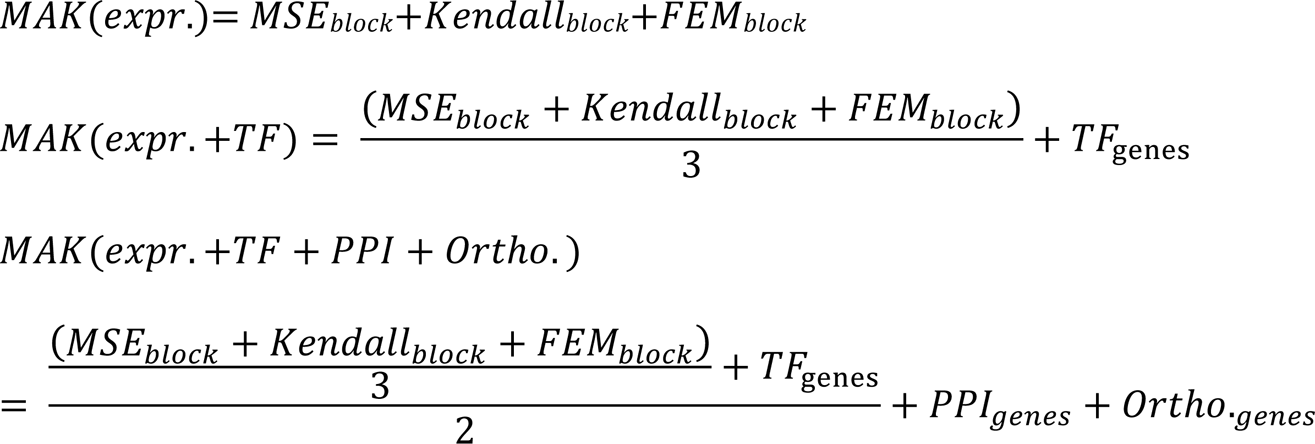

These equations represent the full criterion scoring functions used in this study. The gene expression coherence portion of the functions corresponds to the sum of the mean squared error (MSE) criterion, Kendall’s W statistic criterion, and the fixed effects model (FEM) criterion. This expression coherence criterion was chosen based on simulated data evaluations (Sup. Info.). The other criteria in this equation represent experimental TF binding data (TF_genes_), protein interactions (PPI_genes_), and gene features (Ortho._genes_). In all cases the criteria values used were the corresponding criterion score computed using the null model.

### *De novo* bicluster discovery starting points

To automatically obtain starting points representative of the major data associations present in the data set we use two-dimensional hierarchical clustering (2D-HCL). Since determining the number of expected clusters *a priori* can be computationally expensive and may bias the final results, instead we specify a range of starting point sizes and tile the dataset with exclusive starting points. The number of starting points determines the number of trajectories that need to be generated and thus the computational time for one round of the MAK bicluster discovery. Using the (1- (1+Pearson)/2) correlation as a distance measure for the absolute values of the dataset we hierarchically cluster the data on the first axis using complete linkage (or Euclidean distance and Ward’s criterion). The resulting clustering tree is cut at a height such that no resulting cluster is smaller or larger than a specified size. Next, each of these N clusters from the first axis are clustered with respect to the remaining axis and the resulting second axis clustering trees are similarly cut to respect size range. This procedure gives a set of starting points, starting from the first axis, which tile the entire dataset with blocks obeying size restrictions. The procedure is repeated by clustering on the second axis, cutting according to size, and then clustering each of the M second axis clusters by their first axis, resulting in another complete two-dimensional data clustering and tiling of the input data. We use the union of the first axis and second axis data tilings as the starting points for *de novo* bicluster discovery.

### MAK implementation

The MAK algorithm is implemented in Java, except for most bicluster scoring criteria, the null smoothing, and starting point generation which are implemented in R and accessed using the JRI Java to R interface in the rJava R package (74). The algorithm can be executed using the RunMiner class on a single CPU and if null distributions are pre-generated, this single CPU mode can fully utilize the bicluster criteria parameterization with null distributions. The MAK bicluster discovery pipeline, MAKflow, requires a CPU cluster and is implemented in Java with a limited number of Bash shell (75) and Slurm (76) cluster resource management commands.

### Input *S. cerevisiae* datasets

We used the *S. cerevisiae* gene expression compendium from Reiss et al. (13) consisting of 6160 genes and 667 conditions with row and columns means of values approximately zero (-0.06±0.75, and -0.06±0.77, respectively). Protein-protein interaction data for *S. cerevisiae* was obtained from BioGrid (77) (version 2.0.58, 11/10/09). Experimental TF-gene association rankings were based on ranks of 8mer microarray intensity values from (73). Phylogenetic profiles for 79 complete fungal genomes excluding plasmids were calculated relative to the *S. cerevisiae* genome using genome annotation data available in the MicrobesOnline public MySQL database version on 1/19/12 (67) (RefSeq Release 44, November 2010). All input datasets are available at https://github.com/realmarcin/MAK_results/tree/master/results/yeast/input/input_data.

### MAK *S. cerevisiae* bicluster discovery strategy

A standard MAK bicluster discovery strategy spans seven stages (Figure 2.B). We refer to results of this discovery strategy, i.e. the pooled, nonredundant trajectory endpoints satisfying a score threshold, as the final MAK biclusters. The first stage uses 2D-HCL (described above) to generate starting points for the gene expression data, although any set of submatrices could be used as starting points. Next we ran MAK trajectories using a sequence of two move types: batch moves followed by single moves. We rely on a greedy yet balanced move type sequence, resulting in a guided pseudo-random walk, which cycles through gene addition, condition addition, gene subtraction, and condition subtraction moves. These trajectories were run until none of the four moves in the move sequence are possible that is until no more moves were available to improve the bicluster score. In this work we only used two rounds of searches but multiple rounds are possible using different random seeds and different sets of starting points. Instead, we focused on results from different bicluster pattern types and different input data types.

For each MAK result, all blocks resulting from two rounds of bicluster searches were pooled and biclusters with scores in the top 66th percentile were retained. This score-filtered set of biclusters was then selected for nonredundancy using a 0.25 overlap threshold leading to final sets of MAK biclusters.

### Selecting a nonredundant bicluster set

Biclustering methods vary in the type of bicluster sets they can identify, namely whether or not biclusters can overlap and how much, as well as returning close variations on a single bicluster theme. For interpretation and general downstream analysis purposes it is necessary to work with a representative set of biclusters devoid of redundancy, while preserving biologically relevant bicluster overlaps. To achieve non-redundancy, we selected representative biclusters by measuring pairwise bicluster overlap using cosine similarity, a symmetric overlap measure. The measure is defined by comparing the number of common gene-experiment pairs in biclusters A and B, normalized by the square root product of the gene (G_A_, G_B_) and experiment dimensions (E_A_, E_B_) of the two biclusters:

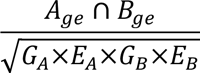

Based on these pairwise overlaps we identified subsets of the bicluster set, where all members overlapped at least 0.25, corresponding to quarter area overlap area assuming equal bicluster sizes. From these overlapping subsets we selected the bicluster with the highest MAK bicluster score. When selecting a nonredundant set from results pooled from multiple bicluster pattern types, the MAK bicluster scores were first scaled to zero mean and unit variance for each pattern type.

### Data-driven bicluster set score for real data in the absence of ground truth

Considering the key properties of a bicluster set, we devised a data-driven bicluster set score consisting of five independent and equally weighted variables. These are i) the maximum of the average of the row or column pairwise bicluster intra-correlations based on the signed or absolute value of the data with mean imputation, ii) the average of the ratio of the bicluster data mean to the overall mean of the dataset, iii) the coverage of conditions by the bicluster set normalized by the number of biclusters, iv) the average number of genes in each bicluster, and v) the average number of conditions in each bicluster. This bicluster set score is designed to include multiple competing features, for example increasing mean bicluster size is expected to lead to decreasing correlation and contrast. On the other hand, increasing correlation and contrast is expected to lead to lower coverage of conditions in the data as well as smaller biclusters.

We removed features with high colinearity (R > 0.5), namely the number of bicluster and gene coverage, resulting in a semi-balanced score with two variables representing the quality of the bicluster patterns (correlation and contrast) with three variables representing the coverage and size of the biclusters (gene coverage per bicluster and mean number of bicluster genes and conditions). Note that this procedure can be repeated for other datasets potentially resulting in a different linearly independent feature set more appropriate for evaluation on that data. Each feature was normalized to the interval [0,1] and the features were combined into a single bicluster set score by calculating the Euclidean distance for each bicluster set relative to the ideal score consisting of maximum values for each feature (1,1,1,1,1).

The data-driven bicluster score does not explicitly take into account the number of biclusters, only indirectly through condition coverage. The true number of biclusters is generally unknown, and depends on the nature of the input dataset, as well as the biclustering approach (i.e., fewer large biclusters or more smaller ones), hence making it a difficult feature to evaluate directly and why we do not model it explicitly in the data-driven bicluster set score. The condition coverage feature also allows controlling for bicluster redundancy since coverage will not increase if the same conditions are repeatedly found in multiple biclusters.

### Supplementary Information and MAK instructions, executable file, examples, source code and results

For additional descriptions of methods see Supplementary Information. The MAK instructions, binary executable (JAR file), examples, and source code are available here https://github.com/realmarcin/MAK. MAK yeast results for different criteria and input data combinations in this study are available here https://github.com/realmarcin/MAK_results/tree/master/results/yeast.

## Acknowledgements

We would like to thank Morgan N. Price, Christopher J. Mungall, Tiffany J. Callahan, Nomi Harris, Michael C. Oldham, Guillaume Cambray, Stefano Cardinale, Michael S. Samoilov, Patrick Flaherty, Maxim Shatsky, and Rachel Brem for discussions and comments on the manuscript. We also thank Keith Keller, Shane R. Cannon, and Shreyas Cholia for systems and HPC advice.

## Supporting Information

**S1 Figure. High throughput evaluation of MAK bicluster criteria on simulated bicluster data.**

Shown are F1 scores and standard deviations (error bars) for MAK criteria with best performance in the five different high throughput biclustering scenarios (constant, row, column, increasing correlation, and overlapping).

**S2 Figure. Evaluation of MAK on the UniBic simulated overlapping bicluster data.** Shown are F1 scores and standard deviations (error bars) for MAK with best performance in the five different high throughput biclustering scenarios (constant, row, column, increasing correlation, and overlapping).

**S1 Table. Biclustering result features and method runtimes.**

Rows are sorted alphabetically. The runtime column represents CPU, GPU, or CPU + CPU cluster total hours. 2D-HCL results are filtered to include only biclusters with mean absolute differential expression > 1.0.

**S2 Table. Size distribution statistics of biclusters from different biclustering methods.** The MAK results are the exemplar with column, constant and row biclusters. 2D-HCL results are filtered to include only biclusters with mean absolute differential expression > 1.0.

**S3 Table. Bicluster functional enrichment across different biclustering methods and MAK results.**

Rows are sorted alphabetically. 2D-HCL results are filtered to include only biclusters with mean absolute differential expression > 1.0.

**S4 Table. Summary of enriched GO terms and term combinations.**

Summary of enriched GO terms and term combinations (TC) found in biclusters from different methods. Total, unique, and missing term combinations as well as individual terms are shown.

**S3 Figure. Comparison of bicluster GO enrichments from different methods on yeast data.**

GO term enrichment was calculated for all results from all methods. For each bicluster, enriched terms were sorted alphabetically, concatenated into a single label, and arranged in a table of combined GO term labels and biclustering methods. Hierarchical clustering was performed on the log2 combined GO term labels. There are relatively few cases of agreement on exact GO term combinations across the different bicluster sets and conversely many methods identify biclusters with unique labels. For 2D-HCL, only biclusters with mean absolute expression > 1.0 were considered. 2D-HCL P-C is with Pearson correlation and complete linkage, while 2D-HCL E-W is with Euclidean distance and Ward’s criterion. Clustered sets of more than 10 GO terms are labeled with the corresponding method or result.

**S1 File. Enriched GO terms and combinations across biclustering methods.**

A list of GO terms and combinations along with methods that reported biclusters with these enrichments. The data is sorted by decreasing count of biclusters enriched for the term or combination.

**S4 Figure. Pairwise bicluster set overlaps from different methods.**

Pairwise bicluster set overlap across biclustering methods and MAK pattern types and data layer combinations. Each cell represents the fraction of biclusters in set A which exhibited fractional overlap > 0.25 with any bicluster in set B. For example, RUniBic results are contained in COALESCE results (A in B) with 1.0 or 100% of biclusters with overlap > 0.25. Conversely, COALESCE results are not contained in RUniBic results (B in A) with 0.0 or 0% biclusters exhibiting overlap > 0.25. This allows to interpret the asymmetrical bicluster set overlap in the context of both sets A and B. 2D-HCL P-C is hierarchical clustering with Pearson correlation and complete linkage, while 2D-HCL E-W is with Euclidean distance and Ward’s criterion; in both cases the biclusters are filtered for absolute mean expression > 1.0.

**S5 Figure. Pairwise bicluster overlap within results from each methods.**

Distributions of bicluster overlap fractions within bicluster sets from each method. For each bicluster in a set, the maximum overlap to any other (non-self) bicluster in the set is recorded and these maximum overlaps are represented as a probability density. Note that MAK biclusters are chosen to maximize the MAK bicluster scoring criterion such that no two biclusters exhibit greater than 0.25 cosine similarity in bicluster gene and condition overlap. The MAK overlap threshold is a user parameter, allowing to control the desired degree of overlap. MAK results are the exemplar MAKcol_const_row(expr.) bicluster set.

**S6 Figure. Pairwise bicluster overlaps between MAK exemplar biclusters and biclusters from all other methods.**

Pairwise comparison of MAK bicluster overlap results with biclusters from other methods. The x-axis represents the maximum coverage fraction of a MAK bicluster from a set of 1655 biclusters from 11 other methods. The y-axis represents the maximum coverage fraction of one of the 1655 biclusters by that MAK bicluster. The red box indicates the set of ‘novel’ biclusters, which have overlap below threshold with values below the diagonal representing novel MAK biclusters and above the diagonal biclusters from the 1655 biclusters from other methods which did not overlap with any MAK bicluster above the 0.25 coverage fraction threshold. The blue box highlights the cases of MAK biclusters covering other biclusters completely; in contrast the maximum coverage of MAK biclusters by biclusters from other methods was 0.73. MAK results are the exemplar MAKcol_const_row(expr.) bicluster set.

**S2 File. Data and biological enrichment features of the 21 novel MAK biclusters.**

Data and biological enrichment features of the 21 novel MAK biclusters compared to 1655 biclusters from 11 top biclustering methods.

